# Mefloquine-induced conformational shift in Cx36 N-terminal helix leading to channel closure mediated by lipid bilayer

**DOI:** 10.1101/2023.12.22.573018

**Authors:** Hwa-Jin Cho, Hyung Ho Lee

## Abstract

Connexin 36 (Cx36) forms interneuronal gap junctions, establishing electrical synapses for rapid synaptic transmission. In disease conditions, inhibiting Cx36 gap junction channels (GJCs) is beneficial, as it prevents abnormal synchronous neuronal firing and apoptotic signal propagation, mitigating seizures and progressive cell death. Here, we present cryo-electron microscopy structures of human Cx36 GJC in complex with known channel inhibitors, such as mefloquine, arachidonic acid, and 1-hexanol. Notably, these inhibitors competitively bind to the binding pocket of the N-terminal helices (NTH), inducing a conformational shift from the pore-lining NTH (PLN) state to the flexible NTH (FN) state. This leads to the obstruction of the channel pore by flat double-layer densities of lipids. These studies elucidate the molecular mechanisms of how Cx36 GJC can be modulated by inhibitors, providing valuable insights into potential therapeutic applications.

## Introduction

Cell-to-cell communication is a crucial process in multicellular organisms, facilitated by intercellular signaling. This communication involves the direct connection of adjacent cells, achieved by the end-to-end docking of two hexameric hemichannels (referred to as connexons) from each cell, resulting in the formation of a dodecameric gap junction channel (GJC)^1^. GJCs facilitate the direct interchange of diverse signals, including electrical impulses and various molecules like ions, second messengers, hormones, and metabolites. Consequently, GJCs play pivotal roles in diverse cellular processes, including neuron electrical synchronization, development, differentiation, and immune response^1,2^. Vertebrate GJCs are formed by connexins, and there are twenty-one human connexin genes.

The proper function of connexins are regulated by multiple factors, including extracellular and intracellular environments, and molecular regulators^1,3^, enabling cells to promptly adjust the turnover rate of GJC at their membrane in response to internal and external signals^4^. Despite the structural homology among connexins, different connexin types exhibit unique properties, including permeability and selectivity^5^. Therefore, comprehending the distinctive attributes of individual connexins and elucidating their regulatory mechanisms across diverse physiological and pathological contexts holds significant importance. However, the precise molecular mechanisms governing how connexins are regulated by intracellular and extracellular factors remain elusive.

Connexin 36 (Cx36) is primarily found in neurons, establishing gap junctions that interconnect neighboring neurons. This molecular architecture leads to the formation of specialized junctions known as electrical synapses, facilitating the rapid transmission of electrical signals and promoting synchronous firing among interconnected neurons. The functionality of Cx36 GJC is influenced by factors such as transjunctional voltage and intracellular pH, cytosolic magnesium ion, arachidonic acid, and regulator molecules like mefloquine (MFQ) and n-alkanols^6–8^. In certain pathological states such as progressive cell death and hypersynchronous neuronal activity, Cx36 GJC participates in the propagation of apoptotic signals and contributes to the occurrence of aberrant synchronous firing, as observed in seizure episodes^9–12^. Thus, the use of inhibitors targeting Cx36 GJC for pharmacological purposes holds promise in potentially disrupting neuronal synchronization during seizure activity and protecting neurons^13,14^, although it is important to recognize the potential for disrupting normal brain function^15,16^. Notably, it has been shown that blocking Cx36 GJC has the capability to efficiently mitigate the severity of seizures, potentially yielding antiepileptic effects^13^. Despite the importance of Cx36 channel inhibition, the mechanism underlying Cx36 channel inhibition remains elusive.

Recently, we resolved the cryo-electron microscopy structures of human Cx36 GJC, unveiling a dynamic equilibrium between its closed and open states^17^. During the process of channel gating, both the N-terminal helices (NTH) and lipid molecules simultaneously play an essential role^17^. In the closed state, the channel pores are filled with two layers of lipids, and the NTHs of Cx36 are dissociated from the pore, forming a flexible NTH (FN) state. Conversely, in the pore-lining NTH (PLN) state, the channel pore is completely open without any obstruction. Given that both PLN and FN states were observed from a single grid sample, an equilibrium of conformational dynamics exists between the closed and open states, which would be modulated by intracellular or extracellular molecules.

Interestingly, small molecules such as MFQ, fatty acids like arachidonic acid, carbenoxolone, and n-alkanol were known to block Cx36 GJC^1,18,19^. Among them, MFQ, an FDA-approved antimalarial compound, is used for the prevention or treatment of mild and moderate malaria^20^, with neuropsychiatric side effects^14,18,21^. Later, it was discovered that MFQ specifically blocks Cx36 GJC, contributing to these side effects, such as spreading depression, NMDAR-mediated neuronal death, tremor, motor deficits, convulsant, stage IV seizures, cell death upon oxygen-glucose deprivation, and hypersensitivity to pain^22–29^. Additionally, it is noteworthy that MFQ exhibits 10- to 100-fold higher sensitivity to Cx36 (IC_50_ value of 310 nM) and Cx50 (IC_50_ value of 1.1 μM) compared to other connexins. Thus, the Cx36-specific channel inhibitor MFQ is regarded as a valuable tool for investigating the regulation mechanism of Cx36 GJC^18^. Another Cx36 GJC inhibitor, arachidonic acid, is an endogenous non-specific channel inhibitor of connexins^7,30^. Under physiological conditions, only a small fraction of Cx36 GJCs, which are assembled into junctional plaques, remains open. However, the number of fractional channels and junctional conductance undergo a significant increase when endogenous arachidonic acid levels are reduced through exposure to fatty acid-free BSA or cytosolic phospholipase A2 inhibitors^7^.

In this study, we elucidate cryo-electron microscopy structures of human Cx36 GJC in complex with its channel modulators like MFQ, arachidonic acid, and 1-hexanol. Notably, these hydrophobic molecules competitively bind to a hydrophobic groove formed by TM1 and TM2, where NTH binds in its open state (PLN state), shifting the conformational dynamic equilibrium of NTHs towards the closed state (FN state). These studies elucidate the previously unresolved molecular mechanisms of how Cx36 GJC can be modulated by inhibitors, providing valuable insights into potential therapeutic applications.

## Results

### Structural features of Cx36 GJC in nanodiscs containing brain lipids

To investigate the distinctive structural characteristics of Cx36 GJC in complex with its inhibitors compared to apo Cx36 GJC, we aimed to reconstitute Cx36 GJC within a lipid nanodisc composed of brain lipids (Cx36_Nano-BL_-WT), considering that Cx36 functions in neuronal GJC. As a first step, we determined the cryo-EM structure of human Cx36 in lipid nanodiscs containing brain lipids (Supplementary Fig. 1). This was necessitated by the absence of an available 3D structure for apo Cx36 GJC reconstructed in brain lipids. The structures of Cx36_Nano-BL_-WT showed both pore-lining N-terminal helix (PLN) and flexible N-terminal helix (FN) states (Supplementary Fig. 1), similar to those solved in a lipid nanodisc containing soybean lipids (Cx36_Nano-SL_-WT, Supplementary Fig. 2b). When overall structures were superimposed, both structures aligned well with identical structural features, including the loose NTH-TM1 loop, the α-to-π-transition of TM1, and the hydrophobic pocket at the channel entrance (Supplementary Fig. 2b, c). However, protomer-focused 3D classification (see method for detail) on Cx36_Nano-BL_-WT revealed that the ratio of Cx36 protomers in the PLN state and the FN state was approximately 3:7 (Supplementary Fig. 3a-b), in contrast to the previously reported 6:4 ratio for Cx36_Nano-SL_-WT^17^. These observations suggest a subtle distinction in the conformational dynamic equilibrium contingent upon the phospholipid composition of the nanodisc.

### The binding of MFQ or arachidonic acid to the Cx36 GJC does not directly obstruct the pore, but induces a conformational change of NTH towards the FN state

Chemical inhibitors, such as mefloquine (MFQ) and arachidonic acid (AA), are widely acknowledged to induce a fully closed state of the Cx36 GJC^1,18^. To explore the molecular mechanism underlying the inhibition of Cx36 GJC by MFQ and AA, we elucidated the cryo-EM structures of Cx36 in complex with MFQ (Cx36_Nano-BL_-MFQ) or AA (Cx36_Nano-BL_-AA) in a lipid nanodisc containing brain lipids at resolutions of 2.7 Å and 2.9 Å, respectively (Fig. 1). Both complex structures exhibited the full FN conformation and displayed similar overall structures to the previously determined Cx36 GJC structures in the FN state, reconstituted in a nanodisc containing soybean lipids (Supplementary Fig. 2b-d)^17^. Notably, both MFQ and AA bind to the hydrophobic groove (H.G.), the highly hydrophobic region within the channel pore (Fig. 1a, b), which is formed by hydrophobic residues from TM1 and TM2 (Fig. 1a, b, upper boxes). While investigating the inhibition mechanism of Cx36 by MFQ and AA, the Cx36-MFQ complex and the Cx32-MFQ complex in detergent micelles were recently reported, showing only the FN state, both in the presence and absence of MFQ^31,32^. This observation aligns with our previously determined structures of human Cx36 in detergents^17^, suggesting that detergents significantly influence the conformation equilibrium between the PLN and FN states.

**Fig. 1.**
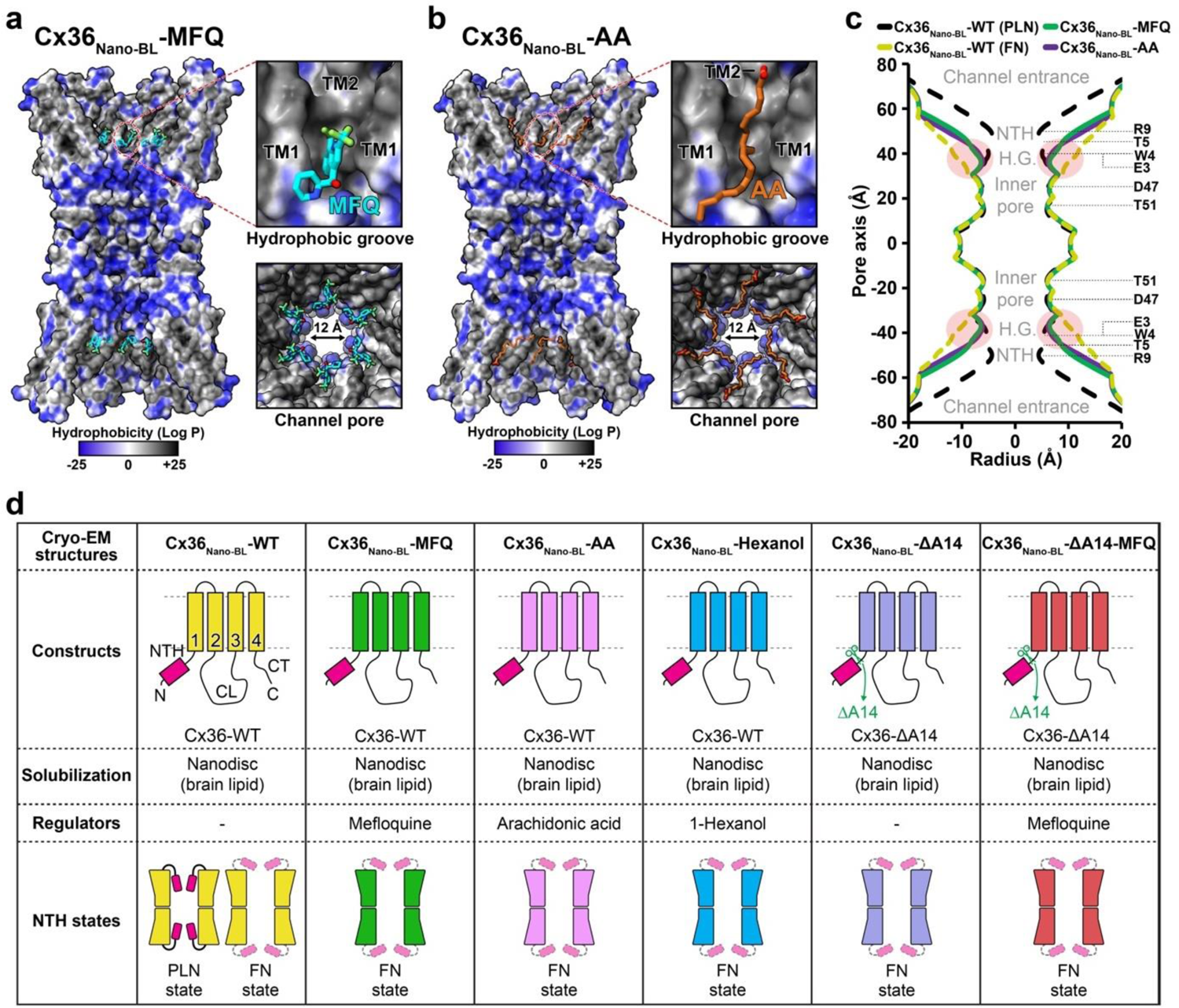
Cryo-EM structures of human Cx36 in complex with mefloquine and arachidonic acid. **a-b**, Cryo-EM maps representing surface hydrophobicity of Cx36_Nano-BL_-MFQ and Cx36_Nano-BL_-AA in cross-sectioned side view. The upper and bottom boxes present close-ups of the hydrophobic groove and a top view, respectively. The water-accessible pore diameter is indicated in the bottom boxes. The surface hydrophobicity (logP) of atomic models is computed and visualized using UCSF ChimeraX^59^, where hydrophobic and hydrophilic surfaces are color-coded in a gradient of dark gray and blue. The channel pore-bound MFQ and AA are depicted in cyan and brown sticks, respectively. **c**, A comparison of the water-accessible pore diameter of Cx36 GJC structures is shown. Cx36_Nano-BL_-WT in PLN and FN states is colored in black and yellow dashed lines, while Cx36_Nano-BL_-MFQ and Cx36_Nano-BL_-AA are colored in green and purple solid lines, respectively. **d**, A schematic diagram summarizing the nine cryo-EM structures of human Cx36 in this study, including nomenclatures used.

When we measured the water-accessible pore diameters of both structures (Cx36_Nano-BL_-MFQ and Cx36_Nano-BL_-AA), they were approximately 12 Å at their narrowest points (Fig. 1a, b, bottom boxes). The binding of MFQ and AA resulted in a reduction of the pore size at the binding site from ∼21 Å to ∼13 Å, indicating a decrease of approximately 8 Å (Fig. 1c, as depicted by the yellow dashed line and the green and purple solid lines). However, their bindings were insufficient to entirely obstruct the channel pore (Fig. 1a, b, bottom boxes). This is in line with the inherent characteristics of the relatively large pore size of Cx36, preventing the direct blockage of the channel pore by the inhibitors. Upon removal of the inhibitors from the atomic models of Cx36_Nano-BL_-MFQ and Cx36_Nano-BL_-AA, the channel pore size closely aligned with that of the previously established Cx36 structure in the FN state (Supplementary Fig. 2d, e). Furthermore, the characteristic α-helix or π-helix structural features observed in previous Cx36 GJC structures in PLN and FN states were also evident at residues 30-33 of TM1 in both Cx36_Nano-BL_-MFQ and Cx36_Nano-BL_-AA (Supplementary Fig. 2c). Taken together, we concluded that the binding of MFQ and AA to the Cx36 GJC does not directly obstruct the pore, but it induces a conformational change of NTH in the FN state, corresponding to the closed state.

### The competitive binding of MFQ or AA to the hydrophobic groove induces the FN state of NTH

We then examined the binding site of MFQ and AA, which is located within the hydrophobic groove of Cx36. This groove is shaped by hydrophobic residues from TM1 (Ile35, Val38 and Ala39) and TM2 (Val80, Ile83, and Ile84) at the interface of the channel entrance and pore (Fig. 2b, c). In the Cx36_Nano-BL_-WT (FN state), weak disordered densities indicative of membrane compounds surround the hydrophobic groove within the channel pore. However, in both Cx36_Nano-BL_-MFQ and Cx36_Nano-BL_-AA, clearly ordered densities corresponding to MFQ or AA emerge (Fig. 2b, c).

**Fig. 2.**
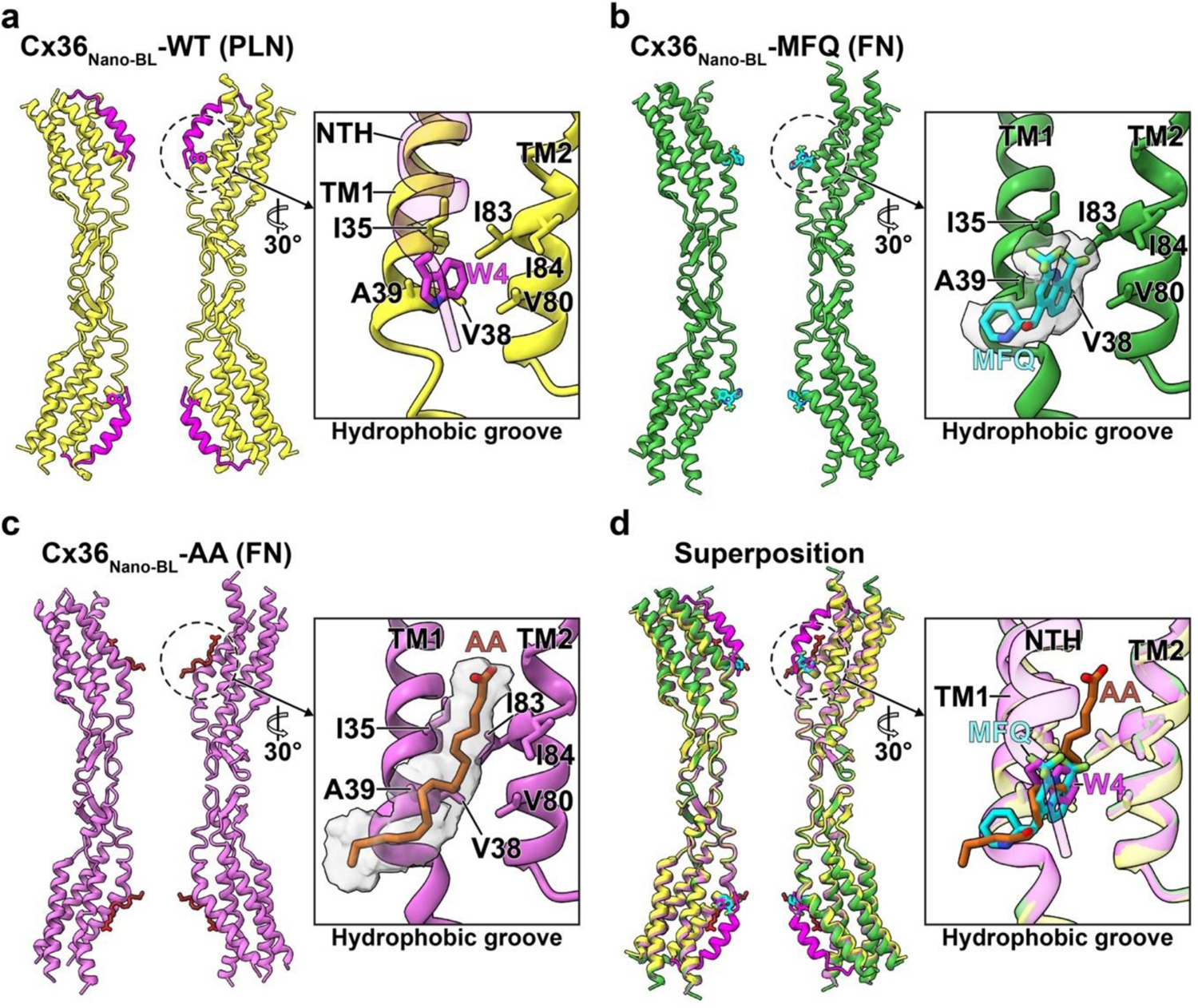
Common hydrophobic binding site of MFQ and AA in Cx36 GJC. **a-d**, The ribbon representation of Cx36_Nano-BL_-WT in PLN (**a**, yellow ribbon), Cx36_Nano-BL_-MFQ (**b**, green ribbon), Cx36_Nano-BL_-AA (**c**, pink ribbon), and their superimposition (**d**). The close-up of the H.G. is presented in boxes. The H.G. is composed of hydrophobic residues from TM1 (Ile35, Val38, and Ala39) and TM2 (Val80, Ile83, and Ile84). The NTH and Trp4 of Cx36_Nano-BL_-WT in PLN are colored in magenta. H.G.-bound MFQ and AA along with their coulomb density maps are shown in cyan, brown, and gray, respectively. H.G.: hydrophobic groove; NTH: N-terminal helix; PLN: pore-lining NTH; FN: flexible NTH; MFQ: mefloquine; AA: arachidonic acid.

Remarkably, both MFQ and AA competitively bind to the H.G. region, the binding pocket of the N-terminal helices. Upon superimposing the structures of Cx36_Nano-BL_-MFQ and Cx36_Nano-BL_-AA onto Cx36_Nano-BL_-WT, Trp4 of NTH in Cx36_Nano-BL_-WT precisely aligns with MFQ and AA of Cx36_Nano-BL_-MFQ and Cx36_Nano-BL_-AA, respectively (Fig. 2d). The binding characteristics suggest that both MFQ and AA bind to the H.G. more effectively than Trp4, i.e., both compounds present broader binding surfaces to the H.G. than Trp4 (Fig. 2), making MFQ and AA more adept at interacting with the H.G. Consequently, the competitive binding of MFQ or AA to the H.G. of Cx36 GJC might prevent Trp4 binding, ultimately triggering the FN state.

To assess the impact of MFQ and AA on the conformational equilibrium of Cx36 GJC, we performed protomer-focused 3D classification following the methodology detailed in our previous work^17^. Strikingly, both inhibitor-bound structures exclusively exhibited FN protomers, with no observation of PLN protomers (Fig. 3 and Supplementary Fig. 3). In the absence of inhibitors, the ratio of PLN to FN protomers was approximately 3:7. However, upon the binding of inhibitors, the ratio completely shifted towards FN protomers (Supplementary Fig. 3). Essentially, the binding of MFQ or AA to the H.G., displacing Trp4, triggers a transition in the conformational dynamic equilibrium towards the FN state. Overall, these findings suggest that MFQ and AA inhibit Cx36 GJC by inducing a conformational shift from the PLN state to the FN state through competitive bindings to the H.G. within the channel pore.

**Fig. 3.**
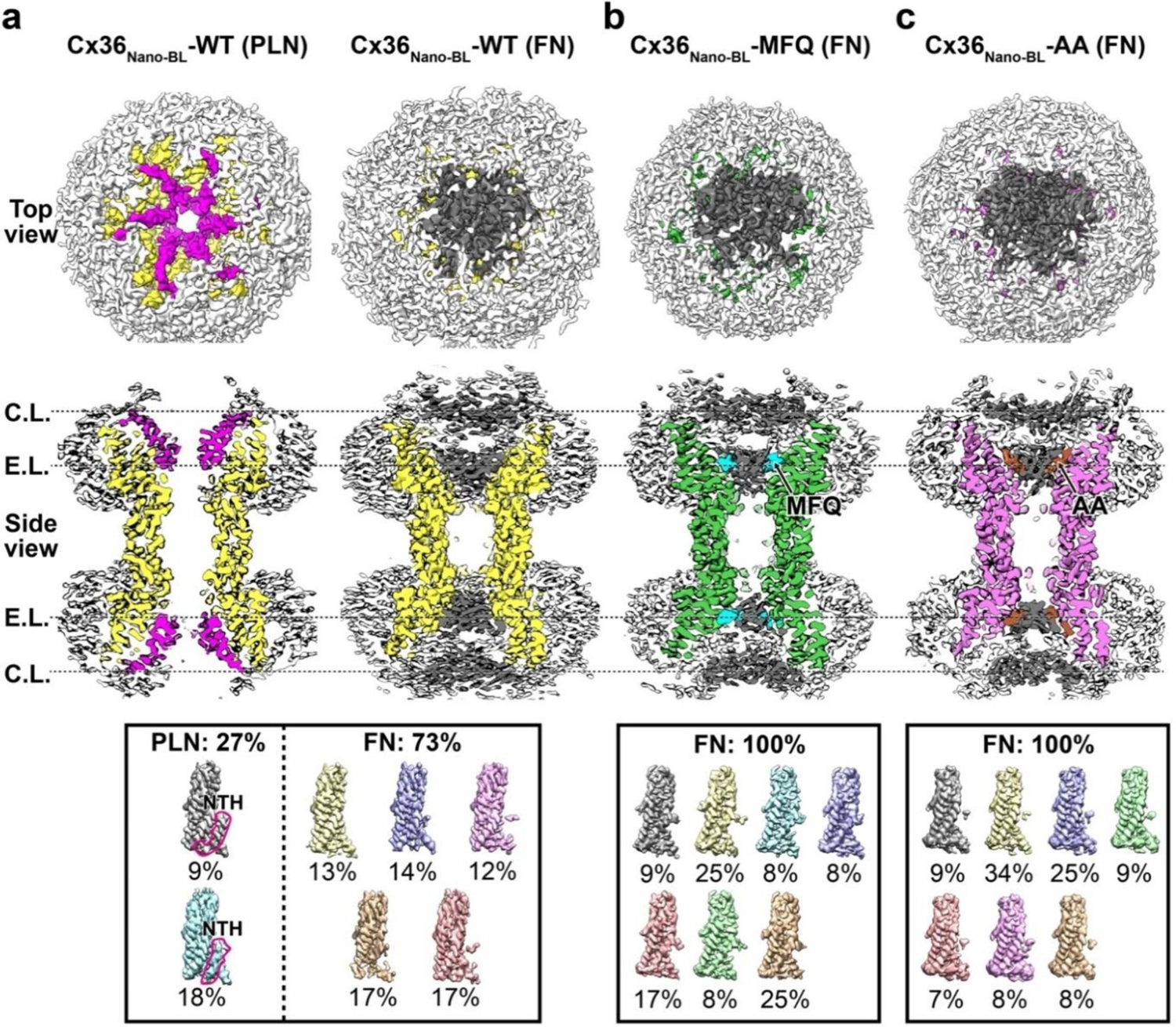
Structural characterization of the pore-occluded states of Cx36_Nano-BL_-MFQ and Cx36_Nano-BL_-AA in lipid nanodiscs. **a-c**, Top and cross-sectioned side views of the cryo-EM reconstruction map with C1 symmetry imposition. The densities of Cx36_Nano-BL_-WT in PLN and FN states (**a**) are displayed in yellow, while Cx36_Nano-BL_-MFQ (**b**) and Cx36_Nano-BL_-AA (**c**) are displayed in green and pink, respectively. The NTHs and double-layered pore-occluding lipids at the C.L. and E.L. are highlighted in magenta and dark gray, respectively. The hydrophobic groove (H.G.)-bound MFQ and AA, as well as lipid nanodiscs, are shown in cyan, brown, and white, respectively. The summary of protomer-focused 3D classification of Cx36_Nano-BL_-WT, Cx36_Nano-BL_-MFQ, and Cx36_Nano-BL_-AA is viewed in the bottom boxes. NTH: N-terminal helix; PLN: pore-lining NTH; FN: flexible NTH; MFQ: mefloquine; AA: arachidonic acid; C.L.: cytoplasmic layer; E.L.: extracellular layer.

### Lipid-mediated occlusion in inhibitor-bound Cx36 GJC structures

In our previous study, we demonstrated a correlation between the NTH conformation (PLN or FN states) and channel gating by lipids^17^. The channel pores of Cx36 GJC are filled with two layers of lipids in the pore-occluded state, with the NTHs dissociated from the pore. Conversely, the channel pore is completely open without any obstruction in the PLN state. This suggests that the binding and dissociation of NTHs to the hydrophobic groove are critical steps in regulating channel opening, although the mechanism by which the conformational equilibrium of NTHs is regulated remains unknown.

Similarly, we observed double-layer density maps in Cx36_Nano-BL_-MFQ and Cx36_Nano-BL_-AA, completely occluding the channel pore (Fig. 3b, c). To avoid artifacts from D6 symmetry imposition during 3D reconstruction, we reconstructed the density maps without imposed symmetry (C1 symmetry). These two layers correspond to the cytoplasmic layer (C.L.) and extracellular layer (E.L.) (Fig. 3). The densities associated with MFQ and AA are located in the E.L. (Fig. 3b, c, cyan and brown densities). While the channel pore was not completely obstructed by the binding of MFQ or AA alone, as demonstrated in Fig. 1, it is fully occluded by the disordered densities (Fig. 3b, c). As elucidated in our prior study, these layers are attributed to lipids, as we excluded the possibility of self-occlusion by components of Cx36 GJC, including NTH, cytoplasmic loop, and C-terminal tail^17^. Moreover, the features of the pore-obstructing densities, including double layers, flatness, and a thickness of ∼4 nm in Cx36_Nano-BL_-MFQ and Cx36_Nano-BL_-AA are highly consistent with those of Innexin-6 hemichannel and pannexin 1 channel^33,34^, suggesting a common mechanism of channel pore occlusion by lipids. Taken together, our structures reveal that the occlusion of the Cx36 channel pore by lipids can be induced by the binding of MFQ or AA.

### Hexanol-driven incomplete occlusion of the channel pore

Next, we hypothesized that a weaker regulator, compared to MFQ and AA, might lead to incomplete occlusion of the Cx36 channel pore, as it could potentially fail to induce a complete shift in the NTH conformation from PLN to FN states. Previous electrophysiological investigations on Cx36 GJC revealed a decrease in channel conductance upon exposure to various alkanols^7^. The n-alkanols are hydrophobic molecules, with hydrophobicity increasing based on the carbon chain length. For instance, 1-hexanol has a hydrophobicity (Log P) of 1.83, while 1-heptanol to 1-decanol exhibit higher hydrophobicity of 2.31, 2.62, 3.06, and 3.44, respectively^35^. Additionally, it is recognized that n-alkanols with different carbon chain lengths have distinct effects on Cx36^35,36^. This observation prompted us to consider that the introduction of such a short alkanol, 1-hexanol, could potentially induce an incomplete shift in the conformational equilibrium of Cx36 GJC towards the PLN state. To test this hypothesis, we determined the cryo-EM structure of Cx36 in the presence of 1-hexanol (Cx36_Nano-BL_-Hexanol) at a resolution of 3.2 Å (Supplementary Fig. 2a).

Indeed, we observed density maps presumed to correspond to 1-hexanol at the hydrophobic groove (Supplementary Fig. 2a), and the binding site of 1-hexanol corresponds to those of MFQ, AA, and W4 of NTH within the H.G. (Supplementary Fig. 2b). When we reconstructed the density maps of Cx36_Nano-BL_-Hexanol without imposed symmetry (C1 symmetry), consistent with Cx36_Nano-BL_-MFQ and Cx36_Nano-BL_-AA, the double-layer densities corresponding to the C.L. and E.L. are significantly weakened, displaying incomplete occlusion (Supplementary Fig. 4c-e). Consequently, we concluded that 1-hexanol, similar to MFQ and AA, can induce the conformational change of NTH toward the FN state through competitive bindings to the H.G. within the channel pore, but it does not result in complete channel occlusion. Taken together, the structural features of Cx36_Nano-BL_-Hexanol suggest that pore occlusion by lipids can be modulated by various inhibitors depending on their capability to remove NTH through competitive bindings to the H.G.

### The preferred binding of MFQ to the H.G. in the PLN state of NTH

We next investigated the potential contribution of the conformational state of NTH to MFQ binding. Since the binding of MFQ to the H.G. is associated with the conformational transition of NTH from PLN to FN states, we postulated that MFQ could bind to the H.G. of Cx36 when NTH is in the PLN state. To test this hypothesis, it was imperative to generate a mutant in which the conformational equilibrium of NTH is entirely shifted to the FN state. During our mutational exploration around the NTH, we uncovered an intriguing variant, Cx36 Ala14-deletion (Cx36_Nano-BL_-ΔA14), demonstrating a complete FN state (Supplementary Fig. 3b). This observation suggested that the Ala14 deletion facilitated the FN state of NTH while the channel pore was obstructed by lipid double layers (Fig. 4c). Although the reason for the impact of Ala14 deletion on the NTH conformation of Cx36 remains unclear, this variant proved valuable in representing a Cx36 with NTH in the full FN state. Subsequently, we determined the cryo-EM structure of the Cx36 Ala14-deletion variant in the presence of MFQ (Cx36_Nano-BL_-ΔA14-MFQ, Supplementary Fig. 6) to assess whether MFQ could bind to the Cx36_Nano-BL_-ΔA14 variant. Remarkably, the MFQ density map disappeared, indicating that MFQ could not bind to the H.G. of Cx36, and thereby supporting the notion that the PLN state of NTH is necessary for MFQ binding (Fig. 4).

**Fig. 4.**
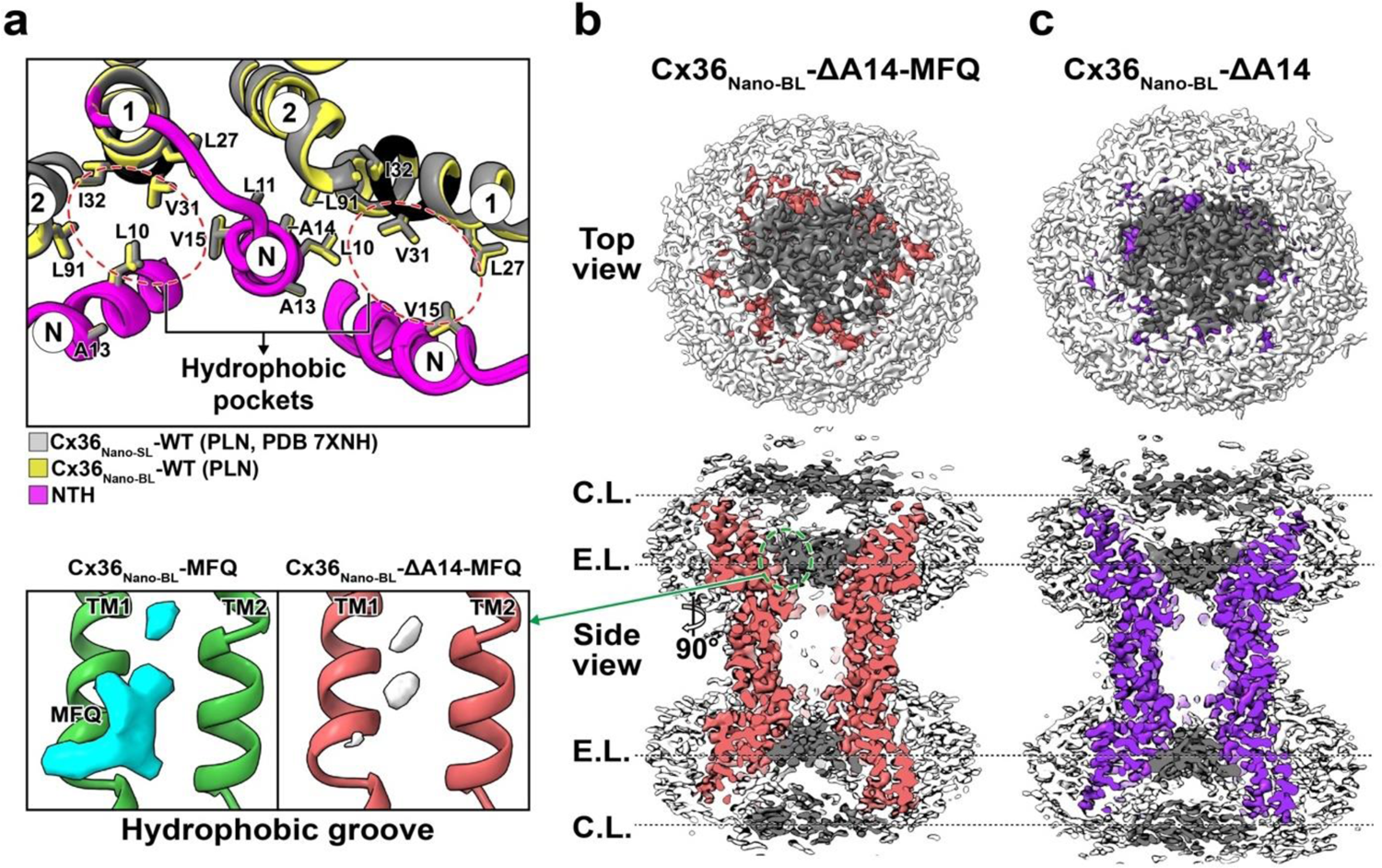
The unfavorable binding of MFQ to the H.G. in the FN state of NTH of Cx36 in lipid nanodiscs. **a**, Structural comparison around the hydrophobic pocket of Cx36_Nano-BL_-WT (yellow) with the PLN state and the previously determined structure of Cx36-WT in soybean lipids (Cx36_Nano-SL_-WT, gray)^17^. The NTH is colored in magenta. The circled N, 1, and 2 denote NTH, TM1, and TM2, respectively. **b-c**, Top and cross-sectioned side views of the cryo-EM reconstruction map with C1 symmetry imposition. The GJC densities of Cx36_Nano-BL_-ΔA14-MFQ (**b**) and Cx36_Nano-BL_-ΔA14 (**c**) are displayed in salmon and purple, respectively. The double-layered pore-occluding lipids at the C.L. and E.L., and lipid nanodiscs are colored dark gray, and white, respectively.

Interestingly, the binding site of MFQ, H.G., is consistent between Cx36 in the PLN state and FN state. Thus, the H.G. seems accessible for MFQ binding in the Cx36_Nano-BL_-ΔA14 variant with the FN state. The binding site of MFQ on Cx36, where Trp4 (or corresponding residues of other connexins) of NTH binds, is highly structurally homologous across other connexins^17,37,38^ (Supplementary Fig. 5b). Hydrophobic residues (Ile35, Val38, Ala39, Val80, Ile83, and Ile84), which interact with MFQ, are well-conserved across connexins as hydrophobic residues (Supplementary Fig. 5a). Thus, the selectivity of MFQ to the H.G. depending on NTH conformation does not stem from the binding site of MFQ. Instead, we reasoned that MFQ may not bind to the H.G. of the Cx36_Nano-BL_-ΔA14 variant with the FN state because the pore is already obstructed by lipid bilayers in the FN state, limiting MFQ’s access to the H.G. (Supplementary Fig. 4g). Collectively, we concluded that the inability of MFQ to bind to the Cx36_Nano-BL_-ΔA14 variant with the fully FN state is attributed to the pore already being obstructed by lipid double layers, preventing MFQ access to the H.G. site. Consequently, the preferential binding of MFQ might occur when it interacts with the H.G. in the PLN state of NTH.

## Discussion

Genetic ablation or pharmacological inhibition of neuronal connexin GJCs has been demonstrated to confer neuroprotection in conditions such as ischemia, traumatic brain injury, glaucoma, and amyotrophic lateral sclerosis^39–41^. Consequently, the pharmacological inhibition of neuronal connexin GJC may offer a strategy for protecting against secondary cell death in such disease contexts. Indeed, several GJC inhibitors, including MFQ, quinine, quinidine, flufenamic acid, 2-APB, and meclofenamic acid, have shown neuroprotective effects by targeting Cx36^22,41–44^. Despite its pharmacological promise and its pivotal role in disease-related neuroprotection, the molecular mechanism of Cx36 channel inhibition remains largely elusive. In this study, we utilized cryo-EM single-particle analysis to investigate the channel inhibition mechanism of human Cx36 GJC, revealing that MFQ, AA, and 1-hexanol bind to the binding pocket of NTH. It induces a conformational change from the PLN state to the FN state, leading to the obstruction of the channel pore by flat double-layer densities of lipids (Fig. 5). Intriguingly, given the inherent characteristics of the relatively large pore size of Cx36, MFQ itself cannot plug the channel pore directly but indirectly induces the conformational changes of NTH, which are coupled with the pore obstruction by lipid double layers (Fig. 3).

**Fig. 5.**
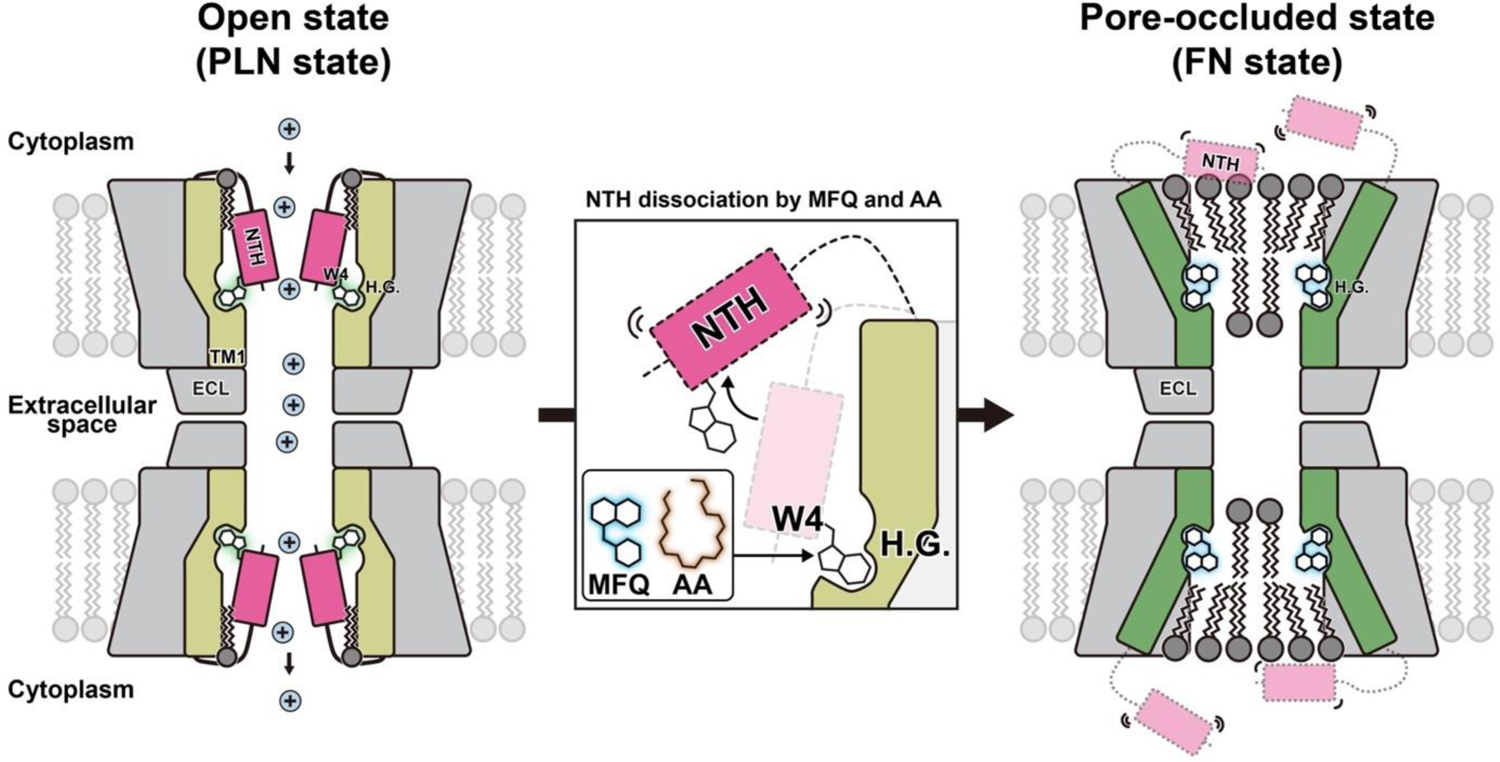
Schematic model for channel inhibition of Cx36 GJC by chemical inhibitors. Schematic representation of the open and pore-occluded states of Cx36 GJC. The NTH, TM1 in the PLN and FN states, TM2-4 and ECL1-2, lipids in the channel pore, lipid bilayer, and cation are colored magenta, yellow, green, gray, dark gray, light gray, and sky blue, respectively. Trp4 of PLN and MFQ are highlighted in green and blue, respectively. MFQ competitively binds to the H.G., where the NTH binds through Trp4. This induces a conformational change from PLN to FN state, shifting the conformational dynamic equilibrium of Cx36 GJC toward the FN state.

In terms of electrophysiology, there are two gating mechanisms for connexin GJCs^45^. One is fast gating, which involves rapid transitions (< 1 ms) between the open state and residual states. The other is slow gating, which involves slower transitions (> 10 ms) between the open state and the fully closed state^45,46^. It is generally considered that the slow gating of connexins is sensitive to factors such as transjunctional voltage, intracellular Ca^2+^, and pH, as well as chemical inhibitors like MFQ and AA^45,46^, where MFQ is classified as an inhibitor that affects the slow gating of Cx36^45,47,48^. In this sense, our proposed mechanism of Cx36 inhibition by MFQ provides evidence of the relationship between the H.G. and slow gating. Indeed, the binding site of MFQ consists of regions that are considered to play a role in slow gating of connexin GJCs, i.e., specific pore-lining residues, including Met34 of Cx26, Leu35, Ala39, Glu43, Gly46, and Asp51 of Cx46, and residues from Phe43 to Asp51 of Cx50, have been identified as contributors to slow gating^45,49–51^. Although a substantial body of evidence is required to fully prove the slow gating mechanism, our study suggests that the lipid-mediated Cx36 blocking by the binding of MFQ to H.G. might be partly related to the slow gating mechanism.

Fatty acids such as arachidonic acids (AA) constitute a component of the cytoplasmic membrane and play a role in regulating the junctional conductance of GJCs. It was reported that AA exhibits inhibitory effects on connexin GJCs, and it is considered to be among the factors contributing to the diminished open probability of Cx36, functioning as an inhibitor^7^. Interestingly, through the structural analysis of Cx36_Nano-BL_-AA, we confirmed that AA binds to the H.G., analogous to the interaction observed with MFQ. Despite difficulties posed by the weak density map, we fitted the carboxyl group of AA facing the cytosol, taking into account the arrangement of adjacent residues. It is intriguing to note that MFQ and AA share the same binding site, H.G., of Cx36, as well as the lipid-mediated blocking, suggesting that MFQ and AA share the same inhibition mechanism. Indeed, similar to the structure of Cx36_Nano-BL_-MFQ, the conformational equilibrium of Cx36_Nano-BL_-AA shifted toward the FN state (Supplementary Fig. 3).

It has been demonstrated that MFQ exhibits higher inhibition sensitivity to Cx36^18^ compared to other connexins. One might expect that the MFQ binding site on Cx36 would be more specific, thus increasing the binding affinity of MFQ. However, the binding site of MFQ on Cx36 shows a notable degree of conservation based on the alignment of amino acid sequences across all connexins (Supplementary Fig. 5a). An alternative hypothesis is that other distinct structural features beyond the H.G. region contribute to the specificity of MFQ to Cx36, such as the loosely packed NTH conformation due to the insertion of Ala14, exclusively observed in Cx36. As a result, among currently available connexin structures in the PLN state, Cx36 exhibits relatively broader hydrophobic pockets that serve as binding sites for a hydrophobic molecule (Supplementary Fig 5c)^17^. The broader hydrophobic pocket is primarily attributed to the longer NTH-TM1 loop (residues Ala13-Ser19) owing to the insertion of Ala14 (Supplementary Fig. 5a). This hydrophobic pocket of Cx36 might contribute to the easier access of MFQ to the H.G., given that the position of the hydrophobic pocket allows it to connect the cytosolic region and the H.G. of Cx36. While this hypothesis could be corroborated through mutational analysis of the hydrophobic pocket, designing mutants poses a challenge since the mutation must not only refrain from impacting the NTH conformation but also solely focus on diminishing the hydrophobicity of the core. This intricacy warrants further investigation in subsequent studies.

Overall, we showed that the hydrophobic groove of Cx36 serves as a common binding site for MFQ, AA, and 1-hexanol. Their interaction triggers a conformational change of NTH from the PLN state to the FN state, consequently shifting the conformational dynamic equilibrium of NTH toward the FN state. Given that the conformational shift from PLN to FN states leads to the obstruction of the channel pore by flat double-layer densities of lipids, we concluded that MFQ, AA, and 1-hexanol share a common inhibition mechanism for Cx36. This implies that the H.G. site of Cx36 might serve as a regulatory site for the NTH conformation, which can be utilized for designing an inhibitor to Cx36. It is tempting to speculate that the NTH conformation of other connexins might also be modulated by the binding of inhibitors to the H.G. site, considering the high structural homology of the H.G. in all connexins.

## Methods

### Expression and purification of Cx36-WT and Cx36-ΔA14 in the baculovirus expression system

All protein expression and purification experiments were conducted following our previously reported method^17^. The complete Cx36 gene was subcloned into pFastBac expression vector (Invitrogen) to generate pFastBac-Cx36-WT. This construct was designed for expression of Cx36 as a fusion protein with C-terminal FLAG tags (consisting of 8 amino acids: DYKDDDDK). This plasmid was engineered to create pFastBac-Cx36-ΔA14 through the deletion of the Ala14 by PCR. The *Escherichia coli* DH10Bac strain (Gibco, cat #10361012) was transformed with pFastBac-Cx36-WT and pFastBac-Cx36-ΔA14 resulting in the production of Cx36-WT and Cx36-ΔA14 bacmid, respectively. These bacmids were then transfected into *Spodoptera frugiperda* (Sf9) cells to generate baculovirus containing the Cx36-WT and Cx36-ΔA14 expression cassette, following the manufacturer’s instructions. Sf9 cells were obtained from ATCC (CRL-1711).

Sf9 cells were grown at 27 °C in suspension within ESF293 medium (Expression Systems), supplemented with 0.06 mg/ml penicillin G and 0.1 mg/ml streptomycin (sigma). At 72 hours post-infection, the cells underwent centrifugation at 500×*g* for 10 min. The membrane was solubilized using buffer A [20 mM HEPES pH 7.5 and 200 mM NaCl], along with 5 mM ethylenediaminetetraacetic acid (EDTA) and protease inhibitors (1 mM phenylmethylsulfonyl fluoride (PMSF), 2 μg/ml leupeptin, 2 μM pepstatin A, and 2 μM aprotinin), supplemented with 1% (w/v) LMNG (Anatrace) for a duration of 2 hours at 4 °C. The insoluble fraction was removed through high-speed centrifugation at 100,000×*g* for 1 hour. Subsequently, the soluble fraction was diluted 2-fold with buffer A, supplemented with 0.01% LMNG, and combined with monoclonal anti-FLAG antibody agarose beads (Wako chemicals, cat #016-22784). This mixture was then incubated with gentle rotation at 4 °C for 5 hours. The resin was settled down within the column and washed three times with 10 column volumes (CVs) of buffer B [20 mM HEPES pH 7.5, 200 mM NaCl, 0.01% LMNG, and 0.001% CHS (Sigma)]. The proteins bound to resin were eluted using buffer B supplemented with 450 μg/ml FLAG peptide (sigma) at 4 °C overnight. The resulting eluates were concentrated and further purified via Superose 6 Increase 10/300 column (cytiva), which was pre-equilibrated with buffer B. Peak fractions were concentrated to ∼6 mg/ml, flash-frozen in liquid nitrogen, and stored at −80 °C for subsequent EM grid preparation.

### Reconstitution of Cx36-WT and Cx36-ΔA14 in lipid nanodiscs containing brain lipids

Purified Cx36 proteins were reconstituted into nanodiscs formed by membrane scaffold protein (MSP1E1) and total brain lipids extract (Avanti) containing phosphatidylcholine, phosphatidylethanolamine, phosphatidylinositol, phosphatidylserine and phosphatidic acid. Lipid cake was prepared using inert gas (N_2_) and resuspended in 5/0.5% (w/v) DDM/CHS, resulting in a clear lipid stock solution at ∼10 mg/ml. The pET28a plasmid carrying the MSP1E1 gene was obtained from Addgene (plasmid #20062). The membrane scaffold protein (MSP1E1) was expressed and purified according to a previously described protocol^52^. The purified Cx36 sample was combined with the total brain lipid extract stock (∼10 mg/ml) in a molar ratio of Cx36 to lipids of 1:100. This mixture was then incubated at 4 °C for 1 h. Subsequently, purified MSP1E1 protein was added, achieving a final molar ratio of Cx36:MSP1E1:lipids of 1:0.5:100. The mixture was subjected to gentle rotation at 4 °C for 30 min. For detergent removal and protein-nanodisc reconstruction, Bio-Beads SM2 resin (Bio-Rad, resin 100 mg) was introduced to the Cx36-lipid-MSP mixture. After 2 hours of gentle rotation, the supernatant was collected. An additional round of detergent removal was carried out using 100 mg Bio-Beads SM2 resin. The mixture was then gently rotated overnight at 4 °C. After removal of insoluble particles through centrifugation at 10,000×*g* for 10 min, the resulting supernatant underwent further purification through size-exclusion chromatography using a Superose 6 Increase 10/300 column pre-equilibrated with buffer A. Fractions containing lipid nanodisc reconstituted Cx36 and Cx36-ΔA14 were concentrated to 3∼5 mg/ml, flash-frozen using liquid nitrogen, and stored at −80 °C for the preparation of EM grid. Protein purity and quality were evaluated through SDS-PAGE (Supplementary Fig. 6).

### Cryo-EM specimen preparation and data collection

The chemical regulators of Cx36, including mefloquine (MFQ), arachidonic acid (AA), and 1-hexanol, were purchased from Sigma. Four microliters of purified Cx36 proteins (4-6 mg/ml) were applied onto a negatively glow-discharged holey carbon grid (Quantifoil R1.2/1.3 Cu 300 mesh). For samples involving Cx36-WT, such as Cx36-WT with MFQ (Cx36-MFQ), arachidonic acid (Cx36-AA), and 1-hexanol (Cx36-Hexanol), reconstituted in lipid nanodisc containing brain lipids (Cx36_Nano-BL_-WT, Cx36_Nano-BL_-MFQ, Cx36_Nano-BL_-AA, Cx36_Nano-BL_-Hexanol, respectively), phenylalanine was added at the final concentration of 50 mM to enhance orientation diversity. The grid was blotted and rapidly frozen in liquid ethane utilizing Vitrobot Mark IV (Thermo Fisher Scientific, USA) under conditions of 4 °C and 100% humidity. Cryo-EM images of Cx36_Nano-BL_-WT and Cx36_Nano-BL_-MFQ were acquired at PNU CORE research facilities (Pusan National University, Korea) and Institute for Basic Science (IBS, Korea) using Krios G4 (TFS, USA) equipped with a BioQuantum K3 detector (Gatan Inc, USA). Additionally, Cryo-EM images of Cx36_Nano-BL_-AA and Cx36_Nano-BL_-Hexanol were collected at the Center for Macromolecular and Cell Imaging (Seoul National University, Korea) using Glacios (TFS, USA) equipped with a Falcon 4 detector (TFS, USA). Automated data acquisition was executed in electron counting mode through the use of EPU software (TFS, USA).

### Image processing and reconstruction

The cryo-EM image processing was conducted using CryoSPARC version 4.1.2 or Relion 4.0 software tools (Supplementary Fig. 1)^53,54^. For the Cx36_Nano-BL_-WT dataset, CryoSPARC (v.4.1.2.) was used. Initial steps included patch-based pre-processing, including patch motion correction & patch CTF estimation, on a dataset containing 8,777 movies. Following this, 4,506,088 particles, identified through reference-based auto-picking, were extracted into 300-pixel boxes and then binned by 2. Additional particles with side-view orientations, identified using Topaz, were also extracted into 300-pixel boxes and binned by 2. After three rounds of 2D classification and two rounds of heterogeneous refinement, the well-classified particles were re-extracted into 300-pixel boxes and subjected to another round of 2D classification. A total 201,311 particles were selected for 3D refinement with D6 symmetry, producing a consensus EM density map at a resolution of 2.6 Å. Subsequently, the particles were transferred to Relion for further classification into the PLN state or FN state using the focused 3D classification method previously used in our study^17^. For 3D skip-alignment classification, 201,311 GJC particles with D6 symmetry imposition were utilized. The process involved 20 initial iterations with a regularization parameter (T) of 20, followed by 10 subsequent iterations with an increased T value (T=40). The GJC mask covering GJC in both FN and PLN state was applied during 3D classification (Supplementary Fig. 1). The resulting four classes displayed distinct features: class 2, containing ∼12% of GJC particles, exhibited clear map densities of PLNs, whereas NTH densities remained less defined in class 1, 3, and 4, comprising ∼88% of the particles. To enhance map quality, particles separated into two GJC conformations underwent separate 3D refinement procedures with D6 symmetry imposition and local angular searches. After sharpening, the result showed a 2.5 Å GJC map with PLN state and a 2.9 Å GJC map with FN state. Finally, 24,000 particles were used for the generation of EM density map of Cx36_Nano-BL_-WT in PLN state with D6 symmetry imposition at 2.9 Å and 177,311 particles were used for the generation of EM density map of Cx36_Nano-BL_-WT in FN state with D6 symmetry at the resolution of 2.9 Å.

The processing of the Cx36_Nano-BL_-MFQ dataset was executed using CryoSPARC (v.4.1.2.). To begin, patch-based pre-processing, including patch motion correction and patch CTF estimation, was applied to the dataset, which comprised 6,483 movies. Subsequently, 3,760,541 particles, selected via reference-based auto-picking, were extracted into 340-pixel boxes and binned by 2. Additional particles featuring side-view orientation, identified through Topaz, were also extracted into 340-pixel boxes and binned by 2. Following three rounds of 2D classification and two rounds of heterogeneous refinement, well-classified particles underwent re-extraction into 340-pixel boxes and were subjected to another round of 2D classification. A total 114,504 particles were used for 3D refinement with D6 symmetry, leading to the generation of a consensus EM density map with a resolution of 2.7 Å. These particles were then transferred to Relion and underwent a single round of 3D classification (K=4, T=32). Finally, 30,596 particles were used to generate the EM density map of Cx36_Nano-BL_-MFQ in FN state, with D6 symmetry imposition at a resolution of 2.7 Å (Supplementary Fig. 7).

Similarly, the Cx36_Nano-BL_-AA dataset was processed using CryoSPARC (v.4.1.2.). Patch-based pre-processing was executed on a dataset containing 4,380 movies, with 1,627,296 particles selected for extraction into 420-pixel boxes and binned by 2. Supplementary particles with side-view orientation were also extracted into 420-pixel boxes and binned by 2. After three rounds of 2D classification and three rounds of heterogeneous refinement, selected particles underwent re-extraction into 460-pixel boxes and subjected to one round of 2D classification. A total of 80,129 particles were then employed for 3D refinement with D6 symmetry, resulting in a consensus EM density map with a resolution of 3.4 Å. Transferring the particles to Relion, one round of 3D classification (K=4, T=20) was performed, leading to the selection of 68,153 particles for generating EM density map of Cx36_Nano-BL_-AA in FN state, with D6 symmetry imposition and a resolution of 3.0 Å. As in previous cases, no PLN state was observed during the image processing procedure.

For the Cx36_Nano-BL_-Hexanol dataset, CryoSPARC (v.4.1.2.) was also utilized. Patch-based pre-processing was conducted on a dataset containing 2,400 movies, and a total of 1,354,338 particles were picked through reference-based auto-picking. These particles were extracted into 220-pixel boxes and binned by 2, with additional particles in side view orientation also were extracted and binned by 2. After three rounds of 2D classification and one round of heterogeneous refinement, well-classified particles underwent re-extraction into 240-pixel boxes and were subjected to one round of 3D classification. Utilizing 289,951 particles, a 3D refinement with D6 symmetry was performed, leading to the generation of a consensus EM density map at a resolution of 4.6 Å. After transferring these particles to Relion, one round of 3D classification (K=4, T=20) was performed. Subsequently, 29,372 particles were chosen to generate the EM density map of Cx36_Nano-BL_-Hexanol in the FN state, with D6 symmetry imposition at a resolution of 3.2 Å. As with previous cases, no PLN state was observed throughout the image processing procedure.

The processing of the Cx36_Nano-BL_-ΔA14 dataset was executed using CryoSPARC (v.4.1.2.). To begin, patch-based pre-processing, including patch motion correction and patch CTF estimation, was applied to the dataset, which comprised 6,313 movies. Subsequently, 3,692,234 particles, selected via reference-based auto-picking, were extracted into 320-pixel boxes and binned by 2. Additional particles featuring side-view orientation, identified through Topaz, were also extracted into 320-pixel boxes and binned by 2. Following three rounds of 2D classification and two rounds of heterogeneous refinement, well-classified particles underwent re-extraction into 320-pixel boxes and were subjected to another round of 2D classification. A total 161,099 particles were used for 3D refinement with D6 symmetry, leading to the generation of a consensus EM density map with a resolution of 2.6 Å. These particles were then transferred to Relion and underwent a single round of 3D classification (K=4, T=40). Finally, 63,476 particles were used to generate the EM density map of Cx36_Nano-BL_-MFQ in FN state, with D6 symmetry imposition at a resolution of 2.7 Å.

The processing of the Cx36_Nano-BL_-ΔA14-MFQ dataset was executed using CryoSPARC (v.4.1.2.). To begin, patch-based pre-processing, including patch motion correction and patch CTF estimation, was applied to the dataset, which comprised 15,027 movies. Subsequently, 4,341,261 particles, selected via reference-based auto-picking, were extracted into 360-pixel boxes and binned by 2. Additional particles featuring side-view orientation, identified through Topaz, were also extracted into 360-pixel boxes and binned by 2. Following three rounds of 2D classification and two rounds of heterogeneous refinement, well-classified particles underwent re-extraction into 360-pixel boxes and were subjected to another round of 2D classification. A total 379,039 particles were used for 3D refinement with D6 symmetry, leading to the generation of a consensus EM density map with a resolution of 2.5 Å. These particles were then transferred to Relion and underwent a single round of 3D classification (K=4, T=40). Finally, 51,406 particles were used to generate the EM density map of Cx36_Nano-BL_-MFQ in FN state, with D6 symmetry imposition at a resolution of 2.6 Å. The local resolutions for all reconstructed maps were estimated by the local resolution estimation tool in CryoSPARC (Supplementary Fig. 8).

### Protomer-focused 3D classification for the Cx36_Nano-BL_-WT dataset

All protomer-focused 3D classification was performed as previously reported^17^. In the case of the Cx36_Nano-BL_-WT dataset, the 201,311 particles, used for the generation of the 2.6 Å consensus EM density map, were transferred to Relion, and D6 symmetry expansion was performed. All protomer particles were subtracted using a mask covering a single protomer (Supplementary Fig. 3a). The subtracted protomer particles were subjected to focused 3D classification (K=8, T=20) with the protomer mask and without orientation search. In the resulting classes, five classes showed the flexible NTH conformation (FN protomer), and two classes showed the pore-lining NTH conformation (PLN protomer). The same method was applied to the Cx36_Nano-BL_-MFQ, Cx36_Nano-BL_-AA, Cx36_Nano-BL_-Hexanol, Cx36_Nano-BL_-ΔA14 and Cx36_Nano-BL_-ΔA14-MFQ.

### Model building and refinement

All structural models with acyl chains and/or CHS molecules were built using the Coot program^55,56^. None of the models include CL (Ala102-Glu193) or CT (Trp277-Val321), and all models, except for the PLN state of Cx36_Nano-BL_-WT, do not include NTH (Met1-His18) due to weak map density. The previously determined structures of Cx36_LMNG_-BRIL (PDB 7XKT) and Cx36_Nano_-WT in PLN state (PDB 7XNH) were used as reference for atomic modeling of all structures determined in this study. The head groups of lipids could not be modeled due to weak map density. All structures were refined using phenix.real_space_refine in the PHENIX software and visualized using UCSF Chimera^57,58^.

### Data availability

The data that support this study are available from the corresponding authors upon reasonable request. Fourteen cryo-EM density maps have been deposited in Electron Microscopy Data Bank (EMDB) under accession codes EMD-38318 (Cx36_Nano-BL_-WT in PLN state (D6 symmetry)), EMD-38346 (Cx36_Nano-BL_-WT in PLN state (C1 symmetry)), EMD-38319 (Cx36_Nano-BL_-WT in FN state (D6 symmetry)), EMD-38347 (Cx36_Nano-BL_-WT in FN state (C1 symmetry)), EMD-38327 (Cx36_Nano-BL_-MFQ (D6 symmetry)), EMD-38326 (Cx36_Nano-BL_-MFQ (C1 symmetry)), EMD-38320 (Cx36_Nano-BL_-AA (D6 symmetry)), EMD-38321 (Cx36_Nano-BL_-AA (C1 symmetry)), EMD-38322 (Cx36_Nano-BL_-Hexanol (D6 symmetry)), EMD-38323 (Cx36_Nano-BL_-Hexanol (C1 symmetry)), EMD-38345 (Cx36_Nano-BL_-ΔA14-MFQ (D6 symmetry)), EMD-38357 (Cx36_Nano-BL_-ΔA14-MFQ C1 symmetry)), EMD-38344 (Cx36_Nano-BL_-ΔA14 (D6 symmetry)), and EMD-38356 (Cx36_Nano-BL_-ΔA14 (C1 symmetry)). Seven atomic coordinates for models have been deposited at the Protein Data Bank (PDB) under accession codes 8XGD (Cx36_Nano-BL_-WT in PLN state), 8XGE (Cx36_Nano-BL_-WT in FN state), 8XGJ (Cx36_Nano-BL_-MFQ), 8XGF (Cx36_Nano-BL_-AA), 8XGG (Cx36_Nano-BL_-Hexanol), 8XH9 (Cx36_Nano-BL_-ΔA14-MFQ), and 8XH8 (Cx36_Nano-BL_-ΔA14) (Supplementary Table 1-3). The previously published structures used for structural comparison were obtained from the Protein Data Bank under accession codes 7XNH (Cx36), 7XKT (Cx36), 2ZW3 (Cx26), 6L3T (Cx31.3), 7F94 (Cx43), 7JKC (Cx46), and 7JJP (Cx50).

## Acknowledgements

We thank Prof. Jae-Sung Woo (Korea University, South Korea), Prof. Insuk So (Seoul National University, South Korea), and Dr. Jinsung Kim (Seoul National University, South Korea) for extensive discussion. The cryo-EM experiments were performed at the Center for Macromolecular and Cell Imaging of Seoul National University, Korea Basic Science Institute (KBSI), Institute for Basic Science (IBS), Bio Open Innovation Center (BOIC) of Pohang University of Science and Technology, and Institute of Membrane Proteins (IMP). This work was supported by Samsung Science & Technology Foundation and Research (SSTF-BA2101-13).

## Author contributions

H.J.C. and H.H.L. conceived this project. H.J.C. performed all the experiments. H.J.C. and H.H.L. analyzed the data and wrote the manuscript and H.H.L. directed the work.

