## Supplementary Information for "Mefloquine-induced conformational shift in Cx36 N-terminal helix leading to channel closure mediated by lipid bilayer"

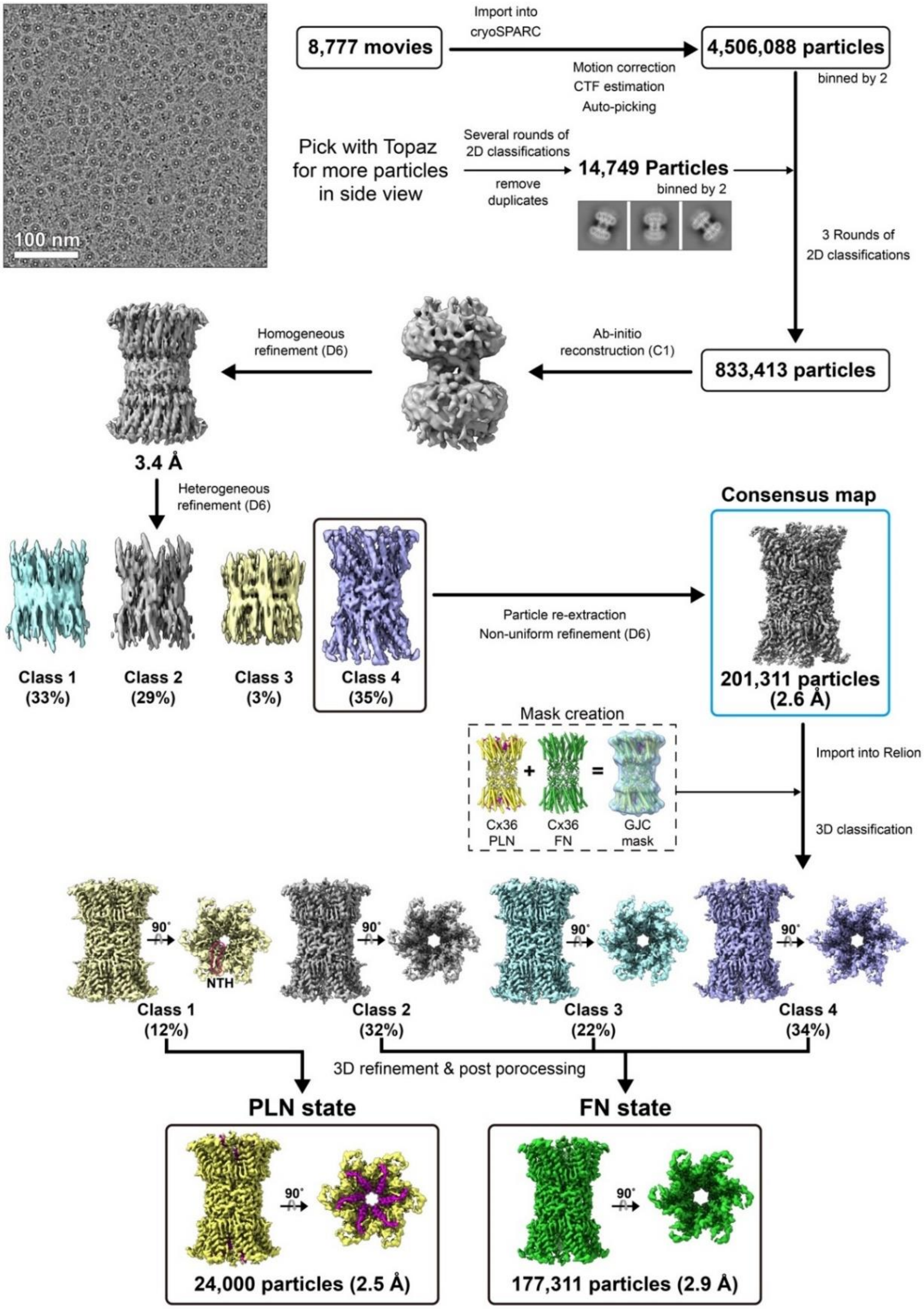


**Supplementary Fig. 1. A flow chart of cryo-EM image processing of $Cx36_{Nano\text{-}BL}$-WT.** The cryo-EM image processing procedure of $Cx36_{Nano\text{-}BL}$-WT is illustrated as a flow chart. This method was also applied to other cryo-EM datasets used in this study.

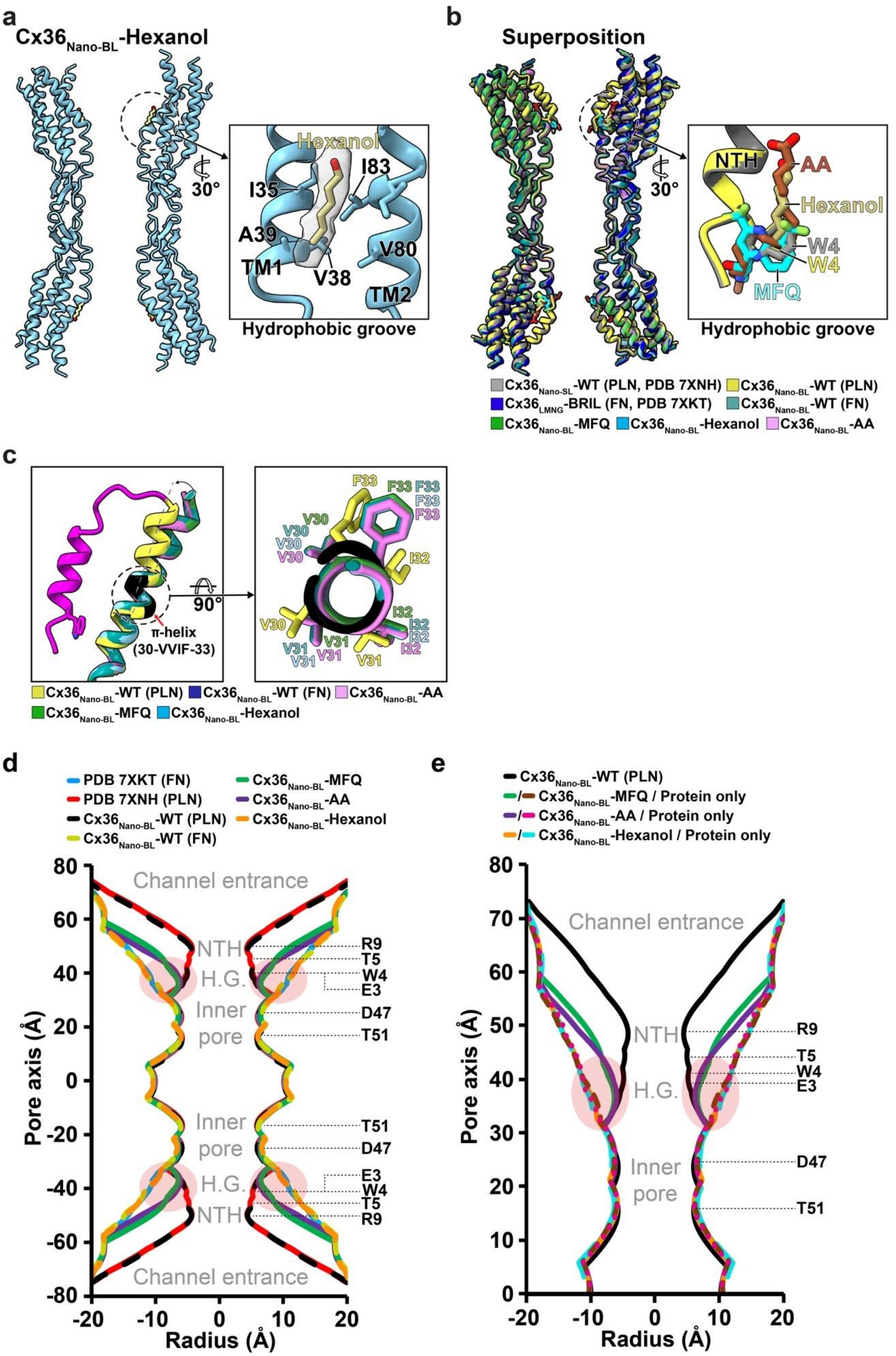

a
Cx36Nano-BL-Hexanol
Hexanol
I35
I83
30°
A39
TM1
V38
V80
TM2
Hydrophobic groove
b
Superposition
NTH
AA
Hexanol
30°
W4
W4
MFQ
Hydrophobic groove
Cx36Nano-SL-WT (PLN, PDB 7XNH)
Cx36Nano-BL-WT (PLN)
Cx36LMNG-BRIL (FN, PDB 7XKT)
Cx36Nano-BL-WT (FN)
Cx36Nano-BL-MFQ
Cx36Nano-BL-Hexanol
Cx36Nano-BL-AA
c
90°
π-helix
(30-VVIF-33)
F33
F33
F33
F33
F33
V30
V30
V30
V30
I32
V30
I32
I32
I32
I32
V31
V31
V31
V31
V31
Cx36Nano-BL-WT (PLN)
Cx36Nano-BL-WT (FN)
Cx36Nano-BL-AA
Cx36Nano-BL-MFQ
Cx36Nano-BL-Hexanol
d
PDB 7XKT (FN)
PDB 7XNH (PLN)
Cx36Nano-BL-WT (PLN)
Cx36Nano-BL-WT (FN)
Cx36Nano-BL-MFQ
Cx36Nano-BL-AA
Cx36Nano-BL-Hexanol
Channel entrance
NTH
H.G.
Inner pore
Inner pore
H.G.
NTH
Channel entrance
R9
T5
W4
E3
D47
T51
T51
D47
E3
W4
T5
R9
Pore axis (Å)
Radius (Å)
e
Cx36Nano-BL-WT (PLN)
/ Cx36Nano-BL-MFQ / Protein only
/ Cx36Nano-BL-AA / Protein only
/ Cx36Nano-BL-Hexanol / Protein only
Channel entrance
NTH
H.G.
Inner pore
R9
T5
W4
E3
D47
T51
Pore axis (Å)
Radius (Å)

**Supplementary Fig. 2. Cryo-EM structure of $Cx36_{Nano\text{-}BL}$-Hexanol and a structure comparion of Cx36 GJC structures with and without inhibitors. a**, The ribbon representation of $Cx36_{Nano\text{-}BL}$-Hexanol along with its Coulomb density map. The close-up of the H.G. is presented in the box. **b**, Structure comparison of previous determined cryo-EM structures of Cx36-WT in the PLN state (PDB 7XNH) and Cx36-WT structures with or without channel regulators. The close-up of the H.G. is presented in the box. **c**, Superposition of N-terminal region to TM1 of Cx36-WT structures with and without channel regulators. The NTH and π-helix around residues 30-VVIF-33 of $Cx36_{Nano\text{-}BL}$-WT in PLN state are colored in magenta and black, respectively. An α-to-π-transition of TM1 is observed during channel opening. **d**, **e**, A comparison of water-accessible pore diameters of Cx36 GJC structures (**d**) and half of protein-only Cx36 GJC structures without regulators (**e**). The previously determined Cx36 structures in the PLN state (PDB 7XNH) and FN state (PDB 7XKT), $Cx36_{Nano\text{-}BL}$-WT in PLN and FN state, $Cx36_{Nano\text{-}BL}$-MFQ, $Cx36_{Nano\text{-}BL}$-AA, $Cx36_{Nano\text{-}BL}$-Hexanol are represented in red, sky blue, black, dark yellow, green, purple, and orange lines, respectively (**d**), and half of manually regulator-removed structures of the $Cx36_{Nano\text{-}BL}$-MFQ, $Cx36_{Nano\text{-}BL}$-AA, and $Cx36_{Nano\text{-}BL}$-Hexanol are shown in brown, magenta, and cyan lines, respectively (**e**). The solvent-accessible pore diameters were calculated using the HOLE program[1].

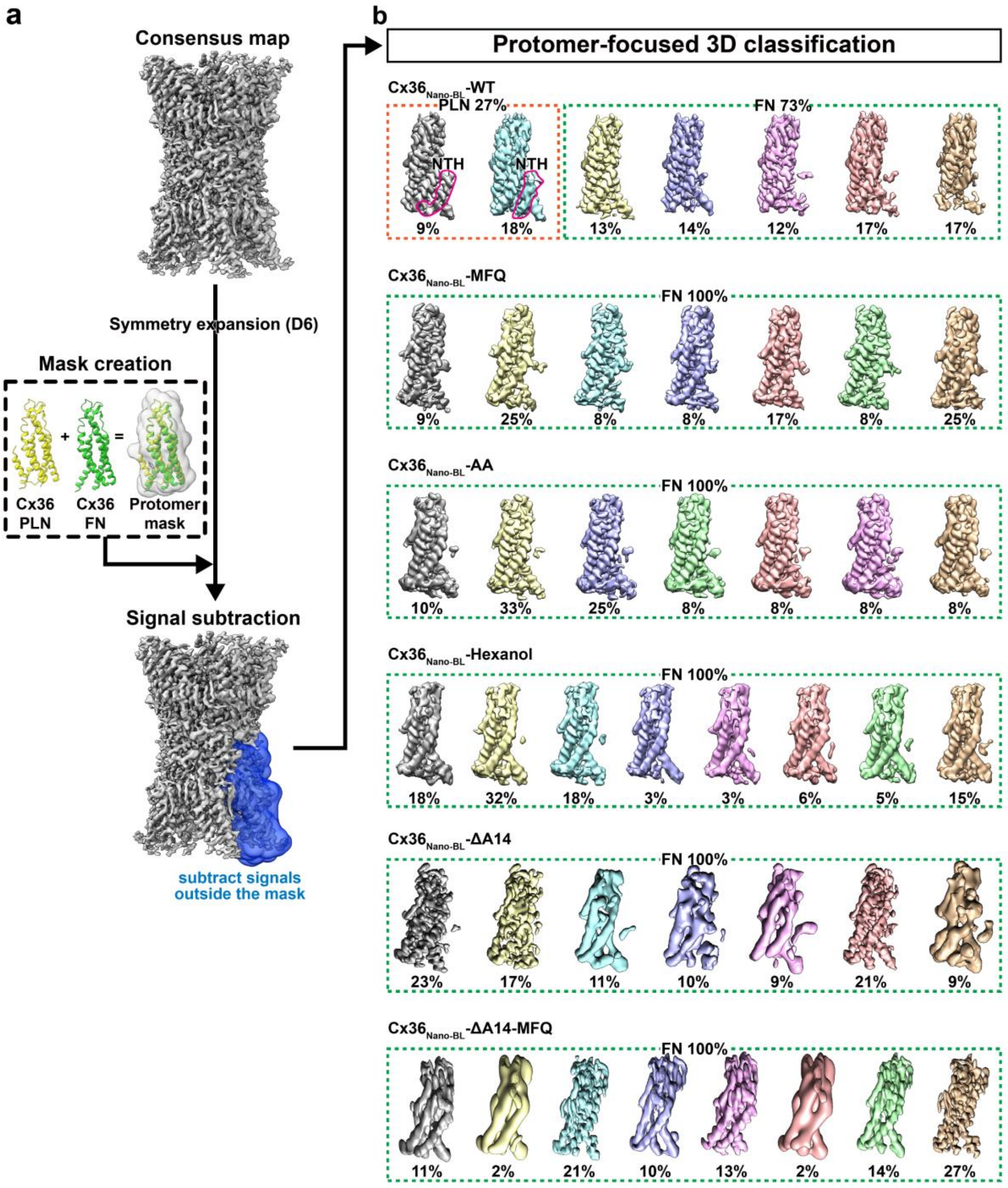


**Supplementary Fig. 3. Protomer-focused 3D classification of Cx36 WT and variants with and without regulators. a**, A simplified representation of the protomer-focused 3D classification with a consensus map (see methods for detail). **b**, 3D classes of protomers. All protomer classes lacking the pore-lining NTH (red solid line) were classified as the FN conformation. The PLN and FN protomer classes are enclosed by orange and green dashed lines, respectively.

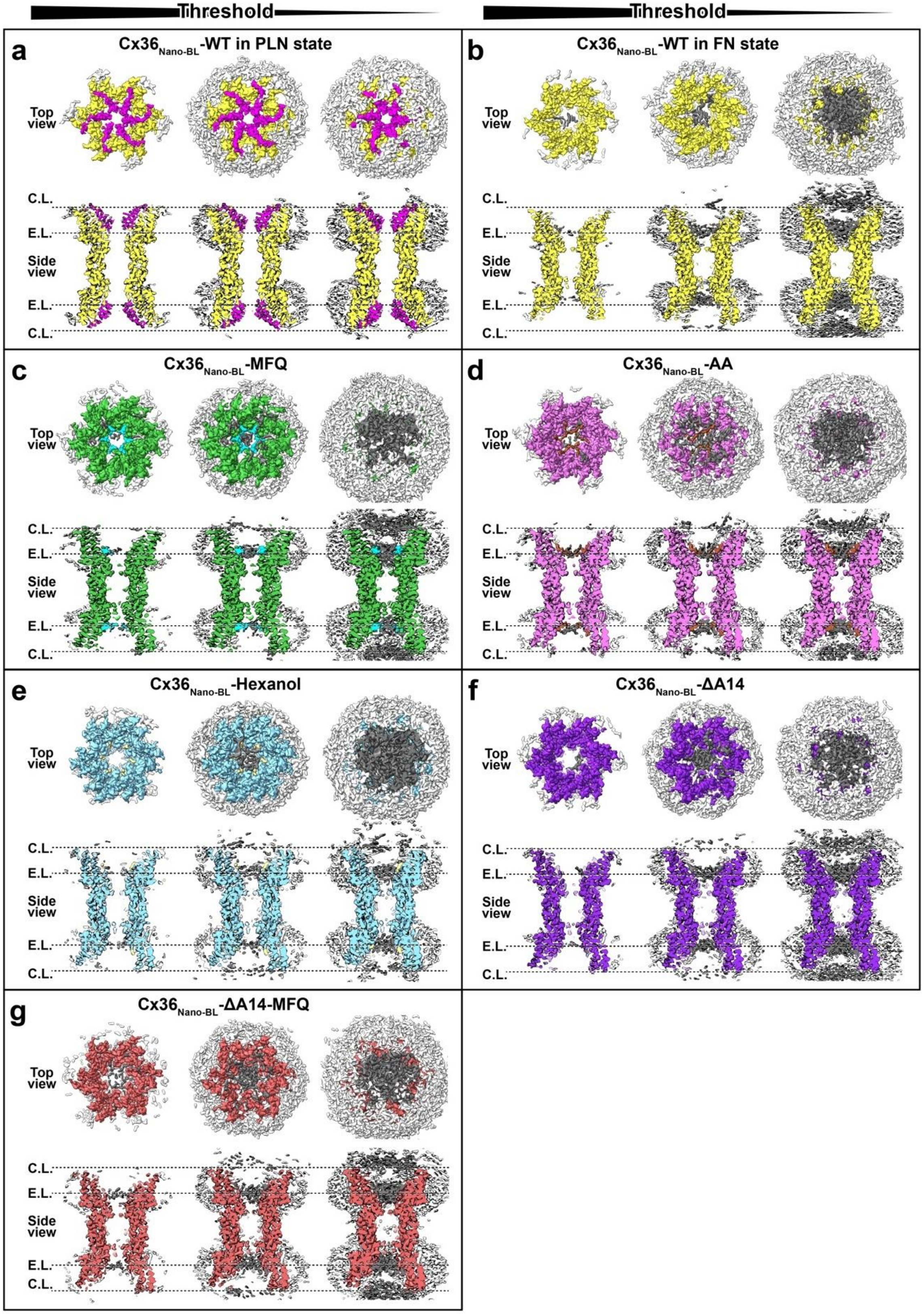

Threshold
Threshold
a
Cx36$_{Nano-BL}$-WT in PLN state
b
Cx36$_{Nano-BL}$-WT in FN state
c
Cx36$_{Nano-BL}$-MFQ
d
Cx36$_{Nano-BL}$-AA
e
Cx36$_{Nano-BL}$-Hexanol
f
Cx36$_{Nano-BL}$-ΔA14
g
Cx36$_{Nano-BL}$-ΔA14-MFQ
Top view
Side view
C.L.
E.L.

**Supplementary Fig. 4. Pore-occluding densities of $Cx36_{Nano\text{-}BL}$-WT and its regulator-bound structures including $Cx36_{Nano\text{-}BL}$-ΔA14 and $Cx36_{Nano\text{-}BL}$-ΔA14-MFQ.** Top views and cross-sectioned side views of cryo-EM map densities with C1 symmetry imposition are displayed at three different map contour levels. The $Cx36_{Nano\text{-}BL}$-WT, $Cx36_{Nano\text{-}BL}$-MFQ, $Cx36_{Nano\text{-}BL}$-AA, $Cx36_{Nano\text{-}BL}$-Hexanol, $Cx36_{Nano\text{-}BL}$-ΔA14, and $Cx36_{Nano\text{-}BL}$-ΔA14-MFQ are colored in yellow, green, orchid, sky blue, purple, and dark red, respectively. The densities of lipid nanodiscs, NTH, MFQ, AA, 1-hexanol, and two pore-occluding lipid layers (C.L. and E.L.) are shown in white, magenta, cyan, brown, khaki, and dark gray, respectively.

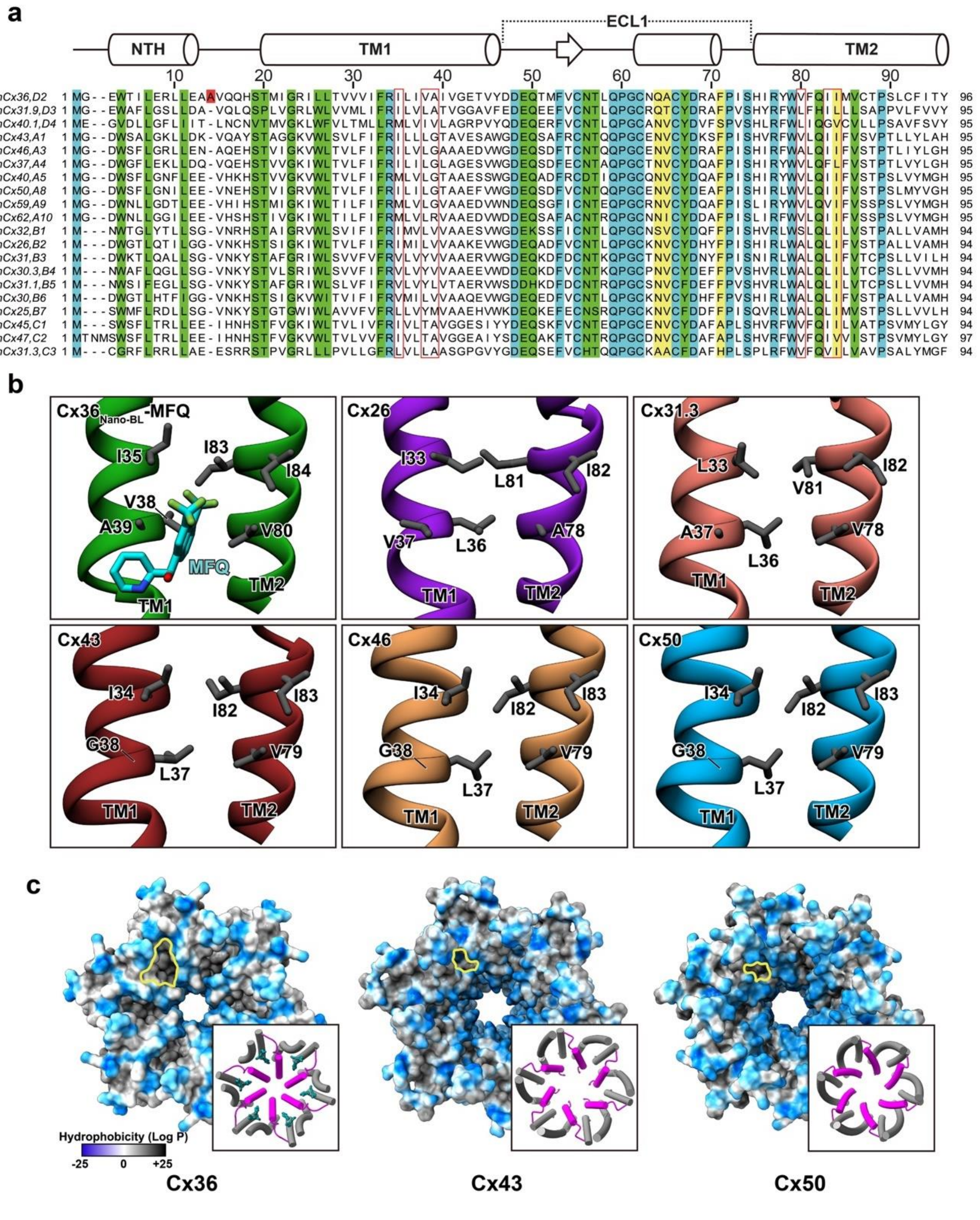


**Supplementary Fig. 5. Amino acid sequence alignment of human connexins and structural comparison of the hydrophobic groove between Cx36 and other connexins. a**, The sequence alignment of all human connexins excluding the highly diversified Cx23. Conserved residues are highlighted in yellow (80%), green (90%), or cyan (100%) colors. Ala14 of Cx36 is shaded in red.

The hydrophobic residues comprising the H.G. of Cx36 and the corresponding residues in other connexins are indicated by red solid lines. **b**, Structural comparison of the H.G. between $Cx36_{Nano\text{-}BL}$-WT-MFQ and currently available other connexin structures, including Cx26 (PDB 2ZW3)[2], Cx31.3 (PDB 6L3T)[3], Cx43 (PDB 7F94)[4], Cx46 (PDB 7JKC), and Cx50 (PDB 7JJP)[5]. Both the amino acids sequence and structure of the H.G. are conserved across all currently available connexin structures. **c**, A top view of $Cx36_{Nano\text{-}BL}$-WT, Cx43 (PDB 7F94), and Cx50 (PDB 7JJP) in PLN state. The surfaces are color-coded based on the hydrophobicity (LogP) of residues, visualized using UCSF ChimeraX software[6]. The cylinder and stub representation of NTH (magenta), TM1 and TM2 (gray) and acyl chain (dark cyan) is shown in boxes.

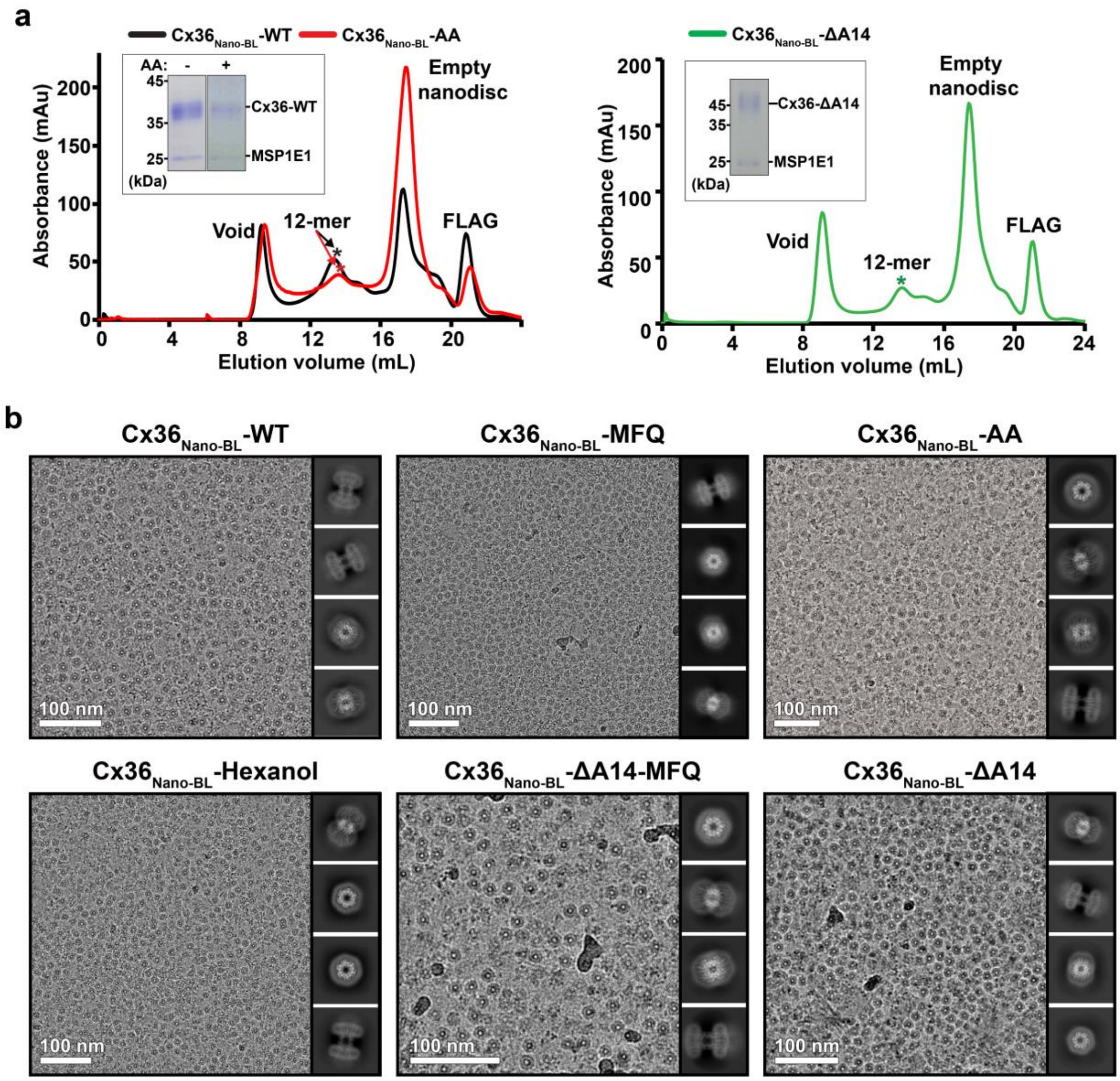


**Supplementary Fig. 6. Purification and cryo-EM imaging of Cx36-WT and Cx36-ΔA14 with and without regulators. a**, Size-exclusion chromatography and SDS-PAGE results of $Cx36_{Nano\text{-}BL}$-WT without AA (black line), and with AA ($Cx36_{Nano\text{-}BL}$-AA, red line), and $Cx36_{Nano\text{-}BL}$-ΔA14 (green line). The SDS-PAGE results of purified $Cx36_{Nano\text{-}BL}$-WT and $Cx36_{Nano\text{-}BL}$-AA, and $Cx36_{Nano\text{-}BL}$-ΔA14 proteins are shown in boxes. **b**, Cryo-EM raw micrographs and 2D classes of $Cx36_{Nano\text{-}BL}$-WT and $Cx36_{Nano\text{-}BL}$-ΔA14 with and without regulators.

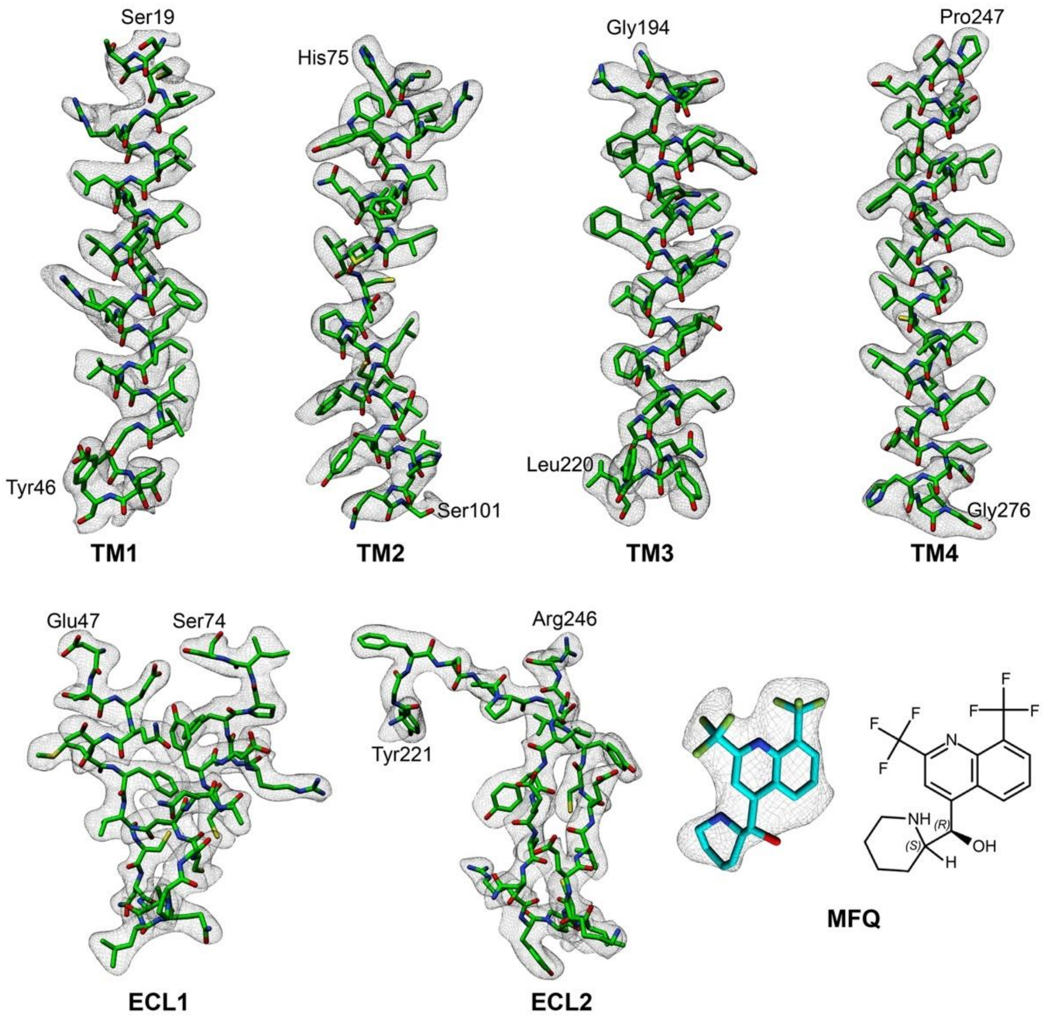


**Supplementary Fig. 7. Cryo-EM map densities of Cx36$_{\text{Nano-BL}}$-MFQ.** Map densities of TM1 (residues 19-46), TM2 (residues 75-101), TM3 (residues 194-220), TM4 (residues 247-276), ECL1 (residues 47-74), ECL2 (residues 221-246), and MFQ are shown.

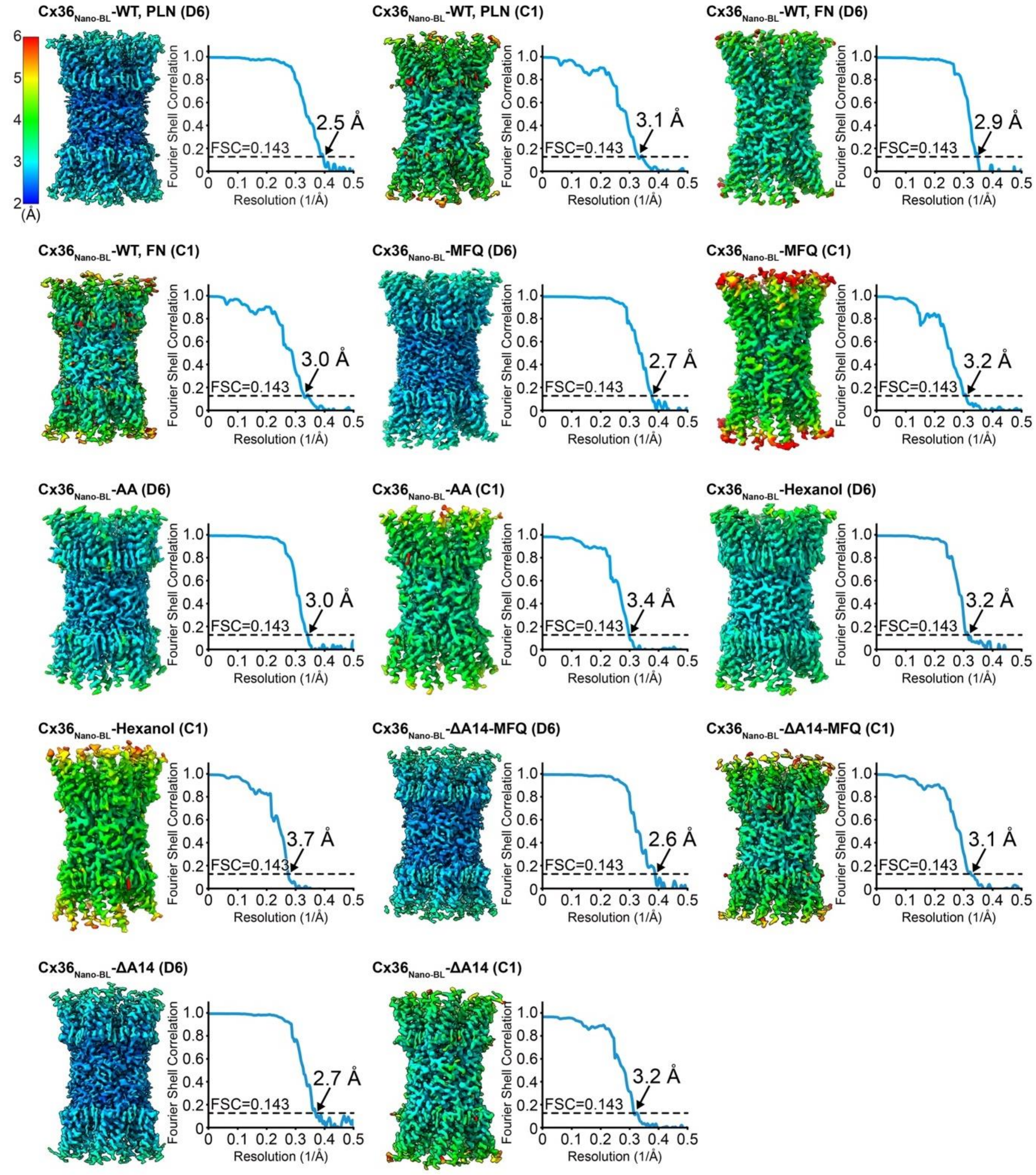


**Supplementary Fig. 8. Local resolution and Fourier shell correlations of fourteen Cx36 maps reconstructed in this study.** The local resolution was estimated using local resolution estimation tool in CryoSPARC and is depicted in a color range from 2 Å (blue) to 6 Å (red). The estimated resolution of the Cx36 structures ranges from 2.5 Å to 3.7 Å. The source data for Fourier shell correlations are provided as a Source Data file.

### Supplementary Table 1. Summarization of cryo-EM structures determined in this study.

| Structure (Map resolution) | PDB ID & EMDB ID | Expression | Environment | Conformation (symmetry) | Protein engineering and additives |
|---|---|---|---|---|---|
| $Cx36_{Nano-BL}$-WT (2.5 Å) | 8XGD & EMD-38318 | Sf9 | Nanodisc (Brain lipid extract) | PLN state (D6) | - |
| $Cx36_{Nano-BL}$-WT (3.1 Å) | N/A & EMD-38346 | Sf9 | Nanodisc (Brain lipid extract) | PLN state (C1) | - |
| $Cx36_{Nano-BL}$-WT (2.9 Å) | 8XGE & EMD-38319 | Sf9 | Nanodisc (Brain lipid extract) | FN state (D6) | - |
| $Cx36_{Nano-BL}$-WT (3.0 Å) | N/A & EMD-38347 | Sf9 | Nanodisc (Brain lipid extract) | FN state (C1) | - |
| $Cx36_{Nano-BL}$-MFQ (2.7 Å) | 8XGJ & EMD-38327 | Sf9 | Nanodisc (Brain lipid extract) | FN state (D6) | Additive: MFQ |
| $Cx36_{Nano-BL}$-MFQ (3.2 Å) | N/A & EMD-38326 | Sf9 | Nanodisc (Brain lipid extract) | FN state (C1) | Additive: MFQ |
| $Cx36_{Nano-BL}$-AA (2.7 Å) | 8XGF & EMD-38320 | Sf9 | Nanodisc (Brain lipid extract) | FN state (D6) | Additive: AA |
| $Cx36_{Nano-BL}$-AA (2.7 Å) | N/A & EMD-38321 | Sf9 | Nanodisc (Brain lipid extract) | FN state (C1) | Additive: AA |
| $Cx36_{Nano-BL}$-Hexanol (3.2 Å) | 8XGG & EMD-38322 | Sf9 | Nanodisc (Porcine brain lipid extract) | FN state (D6) | Additive: 1-Hexanol |
| $Cx36_{Nano-BL}$-Hexanol (3.7 Å) | N/A & EMD-38323 | Sf9 | Nanodisc (Brain lipid extract) | FN state (C1) | Additive: 1-Hexanol |
| $Cx36_{Nano-BL}$-ΔA14 (2.7 Å) | 8XH8 & EMD-38344 | Sf9 | Nanodisc (Brain lipid extract) | FN state (D6) | Deletion: Ala14 |
| $Cx36_{Nano-BL}$-ΔA14 (3.2 Å) | N/A & EMD-38356 | Sf9 | Nanodisc (Brain lipid extract) | FN state (C1) | Deletion: Ala14 |
| $Cx36_{Nano-BL}$-ΔA14-MFQ (2.6 Å) | 8XH9 & EMD-38345 | Sf9 | Nanodisc (Brain lipid extract) | FN state (D6) | Deletion: Ala14<br>Additive: MFQ |
| $Cx36_{Nano-BL}$-ΔA14-MFQ (3.1 Å) | N/A & EMD-38357 | Sf9 | Nanodisc (Porcine brain lipid extract) | FN state (C1) | Deletion: Ala14<br>Additive: MFQ |

**Supplementary Table 2. Cryo-EM data collection, refinement, and validation statistics for D6 symmetry imposed density maps.**

| | **Cx36-WT** | | | | | **Cx36-ΔA14** | |
|---|---|---|---|---|---|---|---|
| | $Cx36_{Nano\text{-}BL}$-WT (PLN state) PDB 8XGD EMD-38318 | $Cx36_{Nano\text{-}BL}$-WT (FN state) PDB 8XGE EMD-38319 | $Cx36_{Nano\text{-}BL}$-MFQ PDB 8XGJ EMD-38327 | $Cx36_{Nano\text{-}BL}$-AA PDB 8XGF EMD-38320 | $Cx36_{Nano\text{-}BL}$-Hexanol PDB 8XGG EMD-38322 | $Cx36_{Nano\text{-}BL}$-ΔA14 PDB 8XH8 EMD-38344 | $Cx36_{Nano\text{-}BL}$-ΔA14-MFQ PDB 8XH9 EMD-38345 |
| **Data collection and processing** | | | | | | | |
| Magnification | 128,000 | 128,000 | 128,000 | 96,000 | 96,000 | 105,000 | 214,000 |
| Voltage (kV) | 300 | 300 | 300 | 200 | 200 | 300 | 300 |
| Electron exposure ($e^{-}/Å^{2}$) | 34.4 | 34.4 | 60.5 | 40 | 40 | 60 | 60 |
| Defocus range (μm) | –0.75 ~ –2.0 | –0.75 ~ –2.0 | –0.75 ~ –2.0 | –1.0 ~ –2.0 | –1.0 ~ –2.0 | –1.0 ~ –2.0 | –0.8 ~ –1.6 |
| Pixel size (Å) | 0.834 | 0.834 | 0.68 | 0.56 | 1.12 | 0.858 | 0.6 |
| Symmetry imposed | D6 | D6 | D6 | D6 | D6 | D6 | D6 |
| Initial particle images (no.) | 2,176,525 | 2,176,525 | 3,760,541 | 1,627,296 | 1,354,338 | 3,692,234 | 4,341,261 |
| Final particle images (no.) | 24,000 | 177,311 | 30,596 | 68,153 | 29,372 | 63,476 | 51,406 |
| Map resolution (Å) | 2.5 | 2.9 | 2.7 | 3.0 | 3.2 | 2.7 | 2.6 |
| FSC threshold | 0.143 | 0.143 | 0.143 | 0.143 | 0.143 | 0.143 | 0.143 |
| Map resolution range (Å) | 2.5 ~ 3.1 | 2.9 ~ 3.6 | 2.7 ~ 3.2 | 3.0 ~ 3.4 | 3.2 ~ 3.6 | 2.7 ~ 3.2 | 2.6 ~ 3.2 |
| **Refinement** | | | | | | | |
| Initial model used (PDB code) | 7XNH | 7XKT | 7XKT | 7XKT | 7XKT | 7XKT | 7XKT |
| Model resolution (Å) | 2.9 | 2.8 | 2.8 | 3.1 | 3.4 | 2.8 | 3.0 |
| FSC threshold | 0.5 | 0.5 | 0.5 | 0.5 | 0.5 | 0.5 | 0.5 |
| Model composition | | | | | | | |
| Non-hydrogen atoms | 18,336 | 16,656 | 17,772 | 17,028 | 16,980 | 16,980 | 18,480 |
| Protein residues | 2,148 | 1,992 | 1,992 | 1,920 | 1,980 | 1,980 | 1,992 |
| Ligands | MC3: 60<br>Y01: 12 | MC3: 48 | MC3: 84<br>YMZ: 12 | MC3: 84<br>ACD: 12 | MC3: 60<br>HE2: 12 | MC3: 72 | MC3: 60 |
| *B* factors ($Å^{2}$) | | | | | | | |
| Protein | 38.08 | 97.98 | 46.12 | 49.47 | 69.72 | 50.71 | 83.44 |
| Ligand | 44.63 | 120.05 | 122.16 | 205.25 | 67.76 | 49.15 | 101.75 |
| R.m.s. deviations | | | | | | | |
| Bond lengths (Å) | 0.003 | 0.002 | 0.003 | 0.002 | 0.003 | 0.003 | 0.004 |
| Bond angles (°) | 0.862 | 0.429 | 0.714 | 0.713 | 0.635 | 0.69 | 0.865 |
| Validation | | | | | | | |
| MolProbity score | 1.51 | 1.25 | 0.79 | 0.87 | 0.98 | 0.93 | 0.93 |
| Clashscore | 9.67 | 4.83 | 1.0 | 1.39 | 2.07 | 1.73 | 1.72 |
| Poor rotamers (%) | 0.21 | 0.2 | 0 | 0.11 | 0.11 | 0.34 | 0.06 |
| Ramachandran plot | | | | | | | |
| Favored (%) | 98.86 | 98.87 | 97.5 | 98.72 | 98.14 | 98.63 | 98.56 |
| Allowed (%) | 1.14 | 1.13 | 2.5 | 1.28 | 1.86 | 1.37 | 1.44 |
| Disallowed (%) | 0 | 0 | 0 | 0 | 0 | 0 | 0 |

### Supplementary Table 3. Cryo-EM data collection and validation statistics for C1 symmetry imposed density maps.

| | **Cx36-WT** | | | | | **Cx36-ΔA14** | |
|---|---|---|---|---|---|---|---|
| | $Cx36_{Nano\text{-}BL}$-WT (PLN state) | $Cx36_{Nano\text{-}BL}$-WT (FN state) | $Cx36_{Nano\text{-}BL}$-MFQ | $Cx36_{Nano\text{-}BL}$-AA | $Cx36_{Nano\text{-}BL}$-Hexanol | $Cx36_{Nano\text{-}BL}$-ΔA14 | $Cx36_{Nano\text{-}BL}$-ΔA14-MFQ |
| | PDB N/A | PDB N/A | PDB N/A | PDB N/A | PDB N/A | PDB N/A | PDB N/A |
| | EMD-38346 | EMD-38347 | EMD-38326 | EMD-38321 | EMD-38323 | EMD-38356 | EMD-38357 |
| **Data collection and processing** | | | | | | | |
| Magnification | 128,000 | 128,000 | 128,000 | 96,000 | 96,000 | 105,000 | 214,000 |
| Voltage (kV) | 300 | 300 | 300 | 200 | 200 | 300 | 300 |
| Electron exposure ($e^-/Å^2$) | 34.4 | 34.4 | 60.5 | 40 | 40 | 60 | 60 |
| Defocus range (μm) | –0.75 ~ –2.0 | –0.75 ~ –2.0 | –0.75 ~ –2.0 | –1.0 ~ –2.0 | –1.0 ~ –2.0 | –1.0 ~ –2.0 | –0.8 ~ –1.6 |
| Pixel size (Å) | 0.834 | 0.834 | 0.68 | 0.56 | 1.12 | 0.858 | 0.6 |
| Symmetry imposed | C1 | C1 | C1 | C1 | C1 | C1 | C1 |
| Initial particle images (no.) | 2,176,525 | 2,176,525 | 3,760,541 | 1,627,296 | 1,354,338 | 3,692,234 | 4,341,261 |
| Final particle images (no.) | 24,000 | 177,311 | 30,596 | 68,153 | 29,372 | 63,476 | 51,406 |
| Map resolution (Å) | 3.1 | 3.0 | 3.2 | 3.4 | 3.6 | 3.2 | 3.1 |
| FSC threshold | 0.143 | 0.143 | 0.143 | 0.143 | 0.143 | 0.143 | 0.143 |
| Map resolution range (Å) | 3.1 ~ 3.8 | 3.1 ~ 3.8 | 3.2 ~ 4.1 | 3.4 ~ 4.0 | 3.6 ~ 4.3 | 3.2 ~ 3.9 | 3.1 ~ 3.8 |
